# The TARANI/ UBIQUITIN SPECIFIC PROTEASE 14 destabilizes the AUX/IAA transcriptional repressors and regulates auxin response in *Arabidopsis thaliana*

**DOI:** 10.1101/850826

**Authors:** Parinita Majumdar, Premananda Karidas, Imran Siddiqi, Utpal Nath

## Abstract

Auxin response is regulated by a group of AUX/IAA transcriptional inhibitors that suppress auxin signaling in the absence of the hormone. While the degradation of these proteins upon auxin signaling has been well studied, the molecular control of their rapid turn-over is not clearly understood. Here, we report that the TARANI/ UBIQUITIN PROTEASE 14 protein in *Arabidopsis thaliana* (Arabidopsis) is required for AUX/IAA degradation. The *tni* mutation was originally identified in a forward genetic screen to isolate mutants with altered leaf shape. Detailed phenotypic analysis revealed that *tni* displays pleiotropic phenotypic alterations that resemble auxin-related defects. The activity of auxin responsive reporters *DR5::GUS*, *DR5::nYFP* and *IAA2::GUS* was reduced in *tni* organs, implying that *TNI* is required for normal auxin response. Genetic interaction studies suggested that *TNI* acts along with *TIR1*, *ARF7*, *AUX1* and *PIN1* – molecules involved in auxin signaling or transport. A map-based cloning approach combined with next-generation sequencing identified *TNI* as *UBIQUITIN SPECIFIC PROTEASE14* which is involved in ubiquitin recycling. In *tni*, the mutant primary transcript is spliced inefficiently, which is predicted to produce an aberrant protein product in addition to the normal protein, where a polypeptide corresponding to the 3^rd^ intron in inserted in-frame within the Zn-finger domain of UBP14. The *tni* plants accumulated poly-ubiquitin chains and excess poly-ubiquitinated proteins due to reduced TNI activity. Improper ubiquitin recycling affected the degradation of DII:VENUS, IAA18:GUS and HS::AXR3-NT:GUS, resulting in their stabilization in the *tni* mutant. Thus, our study identified a function for TNI/UBP14 in regulating auxin response through ubiquitin recycling.

## INTRODUCTION

Coordinated cell division, expansion and differentiation are fundamental to embryonic and post-embryonic organogenesis in plants (ten Hove et al., 2015; Gonzalez et al., 2012; Powell and Lenhard 2012; Sarvepalli and Nath 2018). These cellular processes inturn rely on the turn-over of various regulatory proteins involved in cell-cycle progression (Blilou et al., 2002; Guo et al., 2016), hormonal response (Dharmasiri and Estelle 2004), transcriptional regulation (Osterlund et al., 2000) and gene silencing (Sridhar et al., 2007). Hence, proteostasis is crucialto cellular behavior and depends on the rate of protein synthesis, folding and degradation (Sin and Nollen, 2015). In eukaryotes,the 26S proteasome, a highly selective ATP-dependent protease complex,is the major protein degradation machinery which selectively hydrolyzes cellular proteins tagged with Lys48-linked poly-ubiquitin chains both in the cytosol and nucleus (Hershko and Ciechanover 1992; Thrower et al., 2000). The poly-ubiquitin chains generated upon target protein degradation are hydrolyzed into mono-ubiquitin by a group of processive enzymes known as de-ubiquitinases (DUBs) (Callis, 2014; Yan et al., 2000).These proteases also hydrolyze ubiquitin poly-proteins linked head-to-tail by α-peptide bond and ubiquitin-ribosomal extension proteins into mono-ubiquitin (Callis, 2014). Thus, DUBs are implicated in ubiquitin recycling to accomplish diverse cellular processes.

In Arabidopsis, DUBs are broadly classified into cysteine and metalloproteases based on their catalytic residues (Yan et al., 2000). The domain organization along with the catalytic residues further categorized them into five families:UBIQUITIN-SPECIFIC PROTEASES/ UBIQUITIN BINDING PROTEASES (USPs/UBPs), UBIQUITIN C-TERMINAL HYDROLASES (UCHs), OVARIAN TUMOR PROTEASES (OTUs), MACHADO– JOSEPH DOMAIN (MJD) PROTEASES and JAB1/MPN/MOV34 (JAMM) proteases (Yan et al., 2000; Komander et al., 2009; Isono and Nagel 2014). Among these, the UBPscomprise the largest family with 27 members in Arabidopsis and are further sub-divided into 14 sub-families based on their sequence similarity and domain organization (Yan et al., 2000). The presence of conserved *Cys* and *His* boxes is the hallmark of the UBP proteins. Among 27 *UBPs*, T-DNA knock-out lines *ubp14* and *ubp19* show embryonic lethalityand *ubp15* has narrow and serrated leaves (Liu et al., 2008). The null alleles of the remaining 24 UBPs do not exhibit any discernible phenotypic alteration, suggesting genetic redundancy among them (Liu et al., 2008). However, the higher order mutants of *UBPs* exhibit defects in cell cycle progression, endoduplication, gametogenesis, meristem maintenance and flowering time control (Doelling et al., 2007; Xu et al., 2016, An et al., 2018; Liu et al., 2008).

Most of these UBPs including UBP1-4, UBP12-14 and UBP15 have *in vitro* de-ubiquitination activity against α-linked or iso-linked poly-ubiquitin chainsand ribosomal extension proteins (Isono and Nagel, 2014; March and Farrona, 2018). Apart from ubiquitin recycling, recent studies in Arabidopsis have demonstrated the substrate specificity of DUBs and their pre-eminent role in plant development (Sridhar et al., 2007; Xu et al., 2016; An et al., 2018; Jeong et al., 2017). UBP26 de-ubiquitinates H2B *in vivo*, which is necessary for H3K27me3-mediated gene silencing(Sridhar et al., 2007).Loss-of-function of*UBP26* results in severe defects in seed development due to the loss of repression of *PHERES1*, a MADS box transcription factor (Luo et al., 2008). UBP14 interacts with UVI4, an inhibitor of the anaphase-promoting complex/cyclosome (APC/C) ubiquitin ligase and represses endo-reduplication (Xu et al., 2016). UBP13 de-ubiquitinates RGFR1 and its closest homolog RGFR2, thereby rescuing them from ubiquitin mediated protein degradation by 26S proteasome and thus positively regulating root meristem development through the RGF1– RGFR1–PLT1/2 signaling cascade (An et al., 2018). Similarly, UBP12 and UBP13 de-ubiquitinates poly-ubiquitinated MYC2 *in vitro* and hence act as positive regulators of jasmonic acid (JA) response in Arabidopsis (Jeonget al., 2017). All these recent studies highlight the versatility of DUBs beyond ubiquitin recycling.

Ubiquitin-26S proteasome-mediated turn-over of regulatory proteins is integral to cellular processes ranging from cell cycle progression totranscriptional regulation, hormonal responses and the outcome of biotic/ abiotic stress (Hershko and Ciechanover 1992; Smalle and Vierstra 2004). In Arabidopsis, response pathways to all major phytohormones largely, if not exclusively, rely on the 26S proteasome-mediated protein degradation. The negative regulators of auxin, gibberellic acid (GA) and JA signaling pathways such asAUXIN/INDOLE-3-ACETIC ACID (AUX/IAA), DELLA and JASMONATE-ZIM DOMAIN (JAZ) respectively, undergo poly-ubiquitination and are subsequently degraded by the 26S proteasome. Degradation of these regulatory proteins de-represses the positive regulators of the respective hormone signaling pathways, resulting in the change in gene expression (Daviere and Archard 2013; Gray et al., 1999, 2001; Dharmasiri et al., 2005; Ruegger et al., 1998; Wasternack and Hause 2013).

Among these phytohormones, the role of auxin is well-appreciated as a versatile signaling molecule in organogenesis (Bohn-Courseau, 2010). Auxin regulates a plethora of growth and developmental programs including venation patterning, gravitropism, primary/ lateral root formation and embryo patterning (Hobbie et al, 2000; Swarup et al., 2005; Boerjan et al, 1995; Delarau et al, 1998; Lavenus et al., 2013, ten Hove et al., 2015). Studies over the past few decades have characterized the auxin signal transduction pathway which comprises of TRANSPORT INHIBITOR RECEPTOR1/AUXIN SIGNALING F-BOX Proteins (TIR1/AFBs), AUXIN RESPONSE FACTORs (ARFs) and AUX/IAAs (Ruegger et al., 1998; Gray et al., 1999; Reed 2001; Tiwari et al., 2004). Several genetic and biochemical studies have emphasized on the importance of auxin-dependent degradation of AUX/IAAs by SCF^TIR1/AFBs^ for maintaining the normal auxin response in Arabidopsis (Gray et al, 2001; Leyser et al., 2018). Hence, the auxin level of a given cell is translated into response by activating a set of ARFsthrough release from AUX/IAA repression. Gain-of-function mutations in the degron motif of AUX/IAAs alter their stability, resulting in auxin-related growth defects including venation patterning, lateral root formation, apical dominance and embryo patterning. These mutants include *axr3-1* (Leyser et al., 1996), *axr2-1* (Nagpal et al., 2000), *slr* (Fukaki et al., 2002), *shy2-2* (Tian and Reed 1999), *crane-1*/*iaa18-1*(Uehara et al.,2008; Ploense et al., 2009) and *iaa28-1*(Rogg and Bartel,2001). However, single loss-of-function mutants of *AUX/IAAs* do not exhibit visible phenotypic alterations reflecting a genetic redundancy among themselves (Okushima et al., 2005).

We had earlier reported the isolation and characterization of an Arabidopsis mutant named *tarani* (*tni*)with enlarged, cup-shaped leaves (Premananda Karidas, PhD Thesis, 2014; Karidas et al., 2015). Here, we report the isolation of the *TNI* gene and demonstrate that it encodes UBP14 and is involved in ubiquitin recycling. All the null alleles of *UBP14* reported earlier are embryonic lethal, rendering further studies of this gene in post-embryonic development a difficult task (Doellinget al., 2001; Tzafrir et al., 2002). However, the *tni* allele of *UBP14* is a hypomorph and thus allows us to study the function of this gene in post-embryonic development. The only other reported allele of *UBP14* with post-embryonic viability is*da3-1*which has defective nuclear ploidy and organ growth (Xu et al., 2016). By carrying out detailed phenotypic analysis of *tni* seedlings, we conclude that the TNI/UBP14 protein is required for the optimal auxin response in Arabidopsis. Homozygous *tni* plants showed diverse phenotypic aberrations including defective embryo pattern, tricotyledonous and rootless seedlings, altered tropism of the primary root and fewer lateral roots. These phenotypic defects are also found in mutants with perturbed auxin response. We found thatubiquitin recycling and the turn-over of AUX/IAAswere perturbed in the *tni* seedlings. Taken together, our study demonstrates that the TNI/ UBP14 protein maintains a balance between poly-ubiquitin and mono-ubiquitin which is necessary for the turn-over of AUX/IAAs by 26S proteasome and hence is required for normal auxin response in Arabidopsis.

## RESULTS

### Auxin-relatedpleiotropic phenotypes of *tni*

The *tarani* (*tni*) mutant with altered leaf curvature was originally isolated in a forward genetic screen (Karidas et al., 2015). The mutant leaves showed perturbed pattern of surface expansion and cell division resulting in up-curled lamina as opposed to the flat lamina in wild-type leaves. Detailed phenotypic characterization revealed that *tni* isa pleiotropic mutant with defects in embryonic and post-embryonic development (Fig. 1). During embryogenesis, the*tni* embryo exhibited defects in early cell division pattern (Fig. 1 A-C). In wild-type embryo, the apical cell of the 1-cell pro-embryo undergoes vertical cell division to form the 2-cell pro-embryo and the basal cell undergoes a series of anticlinal division to form the suspensor (Bosca et al., 2011). We observed that the topmost suspensor cell in ∼36% of *tni* embryo underwent periclinal division (N=38) which was never observed in the Col-0 embryo tested (N=30) (Fig. 1A). Besides, while the apical cell of all the Col-0 embryos (N=30)underwentvertical division to form the 2-cell pro-embryo,that in the *tni* embryo underwenthorizontal (∼21%, N=87) or oblique (∼8%, N=87)divisionwith a noticeable frequency (Fig. 1B; Supplemental Fig. S1A and B). Such cell division defects are also observed in the auxin signaling mutant*bodenlos* (*bdl*) and the higher-order polar auxin transport mutant*pin2pin3pin4pin7* (*pin-formed*)(Hamann et al., 1999; Blilou et al., 2005). The hypophysis, a precursor of root stem cell initials, undergoes asymmetric cell division during the dermatogen stage of embryogenesis and forms a lens-shaped cell which is incorporated into the embryo proper at the globular stage (Scheres et al., 1994). A proper lens-shaped cell was observed in all the Col-0 globular embryos (N=30), whereas ∼7% of *tni* embryos(N=95) did not make proper lens-shaped cell (Fig. 1C). The lack of such formative division,which is necessary for specifying root stem cell initials, is expected to result in rootless seedlings. We indeed observed ∼13% rootless seedlings in a *tni* population (N=407), which was never seen in Col-0 (N=127) (Fig. 1D). Perturbed auxin signaling in the gain-of-function mutant *bdl*and in the loss-of-function mutant *monopteros*(*mp*) also result in rootless seedlings (Hamann et al., 1999; 2002).

**Figure 1.** Auxin-related phenotypes in *tni* mutant. **(A)**1-cell pro-embryos. Periclinal cell division defect in the basal cell of 36% *tni* embryo(N=38) indicated by black arrow which was absent in Col-0 (N=30). (B) 2-cell pro-embryos. The apical cell of Col-0 embryo underwent vertical division (N=30), denoted by black arrow whereas horizontal cell division pattern in ∼21% *tni* embryos (N=87) indicated by black arrow. **(C)** Globular stage embryos. In all the Col-0 embryos (N=30), a proper lens shaped cell was formed, magnified in the inset whereas cell division defect was observed in∼7%*tni*embryo (N=95) as shown in the inset. **(D)** Side-view of seedlings. All Col-0 seedlings produced well-formed root system (N=127) while ∼13% *tni* seedlings showed rootless phenotype (N=407).**(E)** 7-day old seedlings. While all Col-0 seedlings produced two cotyledons (N=45), ∼4%*tni* seedlings produced three cotyledons (N=306). **(F)** Heart stage embryos. Col-0 and *tni* embryos showing two and three cotyledon primordia respectively marked in dotted red lines. The meristematic region in Col-0 embryo is false colored in yellow.**(G)** Cleared cotyledons from 7-day old seedlings highlighting venation pattern. Open venation in the *tni*cotyledon is marked by black arrows.**(H)**The percentage of cotyledons showing closed or open venation patterns. N=103 (Col-0) and 90 (*tni*). **(I)** Effect of gravistimulation on the root of 7-day old seedlings. The angles of root bending are indicated by red lines. **(J)** The average angle of curvature of primary roots following gravistimulation. N=52 (Col-0) and 43 (*tni*). Error bars represent SD.The statistical analysis was done by unpaired Student’s *t*-test. *** denotes p<0.0001. The scale bar=5mm.(**K**) 11-day old seedlings highlighting the lateral roots. **(L)**The average number of lateral roots (N=10-13) produced on the primary root of seedlings at indicated days after germination (DAG). Error bars represent SD. Statistical analysis was done by unpaired Student’s *t*-test. *** denotes p<0.0001. The scale bar=5mm. **(M)**Open flowers showing the variation in petal numbers. N=142 (Col-0) and 140 (*tni*).(**N)**Matured seeds showing bigger area in *tni*. The scale bar=5000μm. (O) The average area of seeds. N=15. Error bars represent SD.The statistical analysis was done by unpaired Student’s *t*-test. *** denotes p<0.0001

The post-embryonic developmental defects in *tni* included tricotyledonous seedling,reduced venation complexity in cotyledon with incomplete vein loop,defects in gravitropic response in primary root, fewer lateral roots, larger seeds and increased floral organs (Fig. 1E-O).We estimated that ∼4% of *tni* seedlings (N=306) form three cotyledons(Fig. 1E), while we never observed tricotyledonous Col-0 seedlings in our growth conditions. To address the origin of thisphenotype, we examined the embryos at the transition state from the globular to the heart stage when cotyledon primordia are initiated at the flanks of the embryo apex (ten Hove et al., 2015). Col-0 embryo always formed two cotyledon primordia whereas *tni* embryo occasionally produced three (Fig. 1F). Similar tricotyledonous phenotype is also observed in *pin1* loss-of-function mutant that has perturbed auxin transport (Krecek et al., 2009). Mature *tni* cotyledons showed fewer complete areoles (closed veins) as opposed to four complete areoles produced by the Col-0 cotyledons, suggesting that the complexity of venation pattern was reduced in *tni* (Fig. 1G and H). When a population of 7-dayold seedlings was analyzed for cotyledon vein development, ∼20% Col-0cotyledons (N=103) showed maximal complexity with four complete areoles and the remaining had <4 areoles with some open vasculature (Fig. 1H and Supplemental Fig. 1C and D) (Cnops et al, 2006; Sieburth 1999). By contrast, only ∼10% (N=90) *tni*cotyledons attained maximal complexity with four areoles and another ∼10% exhibited open top vein defect which was never observed in Col-0 (Fig. 1H). Venation defect is also observed in *auxin-resistant mutant6* (*axr6*) and *cotyledon vascular pattern1* (*cvp1*), a mutant with altered auxin distribution(Hobbie et al., 2000; Reinhardt, 2003).

The *tni* mutant exhibited defects in the sub-aerial organs as well (Fig. 1I-L). Primary root in *tni* seedlings showed defects in gravitropic response (Fig. 1I). After gravistimulation, Col-0 primary roots responded to the gravity by bending at an average angle of 83.1° ± 11.50 (N=52) towards gravity. By contrast, the *tni* roots bent only by 56.1° ± 11.0 (N=43)under similar experimental conditions (Fig. 1J), suggesting a reduced response towards gravity in the mutant. Besides altered gravitropic response, *tni* primary roots also produced fewer lateral roots (Fig. 1K andL). In Col-0 seedlings, the number of emerged lateral roots steadily increased from 7 to 11 days after germination (DAG) (Fig. 1L). Though a similar trend was observed in the *tni*seedlings, the number of lateral rootsremained ∼50% lower at all the growth stages measured. Fewer lateral roots have also been reported in mutants with perturbed auxin transport and signaling including *tir1-1*, *auxin1-7* (*aux1-7*) and *auxin response factor7-1* (*arf7-1*) (Ruegger et al., 1998; Marchant et al, 2002; Okusima et al., 2007).

Petal number in *tni*flowers varied from the wild type (Fig. 1M). While all the Col-0 flowers examined (N=142) formed four petals, ∼58% *tni* flowers (N=140) showed 5-6 petals (Fig. 1M). This phenotype is similar to the weaker allele of polar auxin transport mutant *pin1-5* (Yamaguchi et al., 2014).The *tni* plants also produced bigger seeds with nearly 30% increase in seed area compared to wild type (Fig. 1N and O).Bigger seedsare seen in the auxin response mutant *auxin response factor2* (*arf2*), a negative regulator of cell division and expansion in Arabidopsis (Schruff et al., 2005).

A number of mutants in Arabidopsis with perturbed auxin biosynthesis, transport or signaling exhibit aberrant cell division in embryo, patterning defect in cotyledon, rootless seedlings, fewer lateral root, supernumerary petals and altered tropism in root or shoot (Cheng et al., 2007; Swarup et al., 2005; Marchant et al., 2002; Hobbie et al., 2000; Hamann et al., 2002, Ruegger et al., 1998, Bennett et al., 1995). Notably, *tni* exhibited defects in most of these processes suggesting its possible involvement in the auxin pathway.

### Reduced auxin response in *tni*

The auxin-related growth defects of *tni* prompted us to compare auxin response in Col-0 and *tni* seedlings using the auxin-responsive *DR5::GUS*, *DR5::nYFP* and *IAA2::GUS* reporter lines (Ulmosov et al., 1997; Mahonen et al., 2015; Marchant et al., 2002). GUS assay of 3-dayold *DR5::GUS* seedlings showed strongβ-glucuronidase activity throughout the cotyledon margin with the highest signal at the tip, indicative of auxin maxima (Sabatini et al., 1999;Mattsson et al., 2003) (Fig. 2A). No such distinct auxin maxima wasdetected in the *tni* cotyledons, with an overall reduction in GUS signal throughout the cotyledon compared to Col-0. Similar reduction in auxin maximawas also observed at the tip of emerging leaves, primary roots and lateral roots of *DR5::GUStni*seedlings (Fig. 2B-D). Comparison of the fluorescence signal in the *DR5::nYFP*and*DR5::nYFPtni*also detected reduced auxin maxima in the cotyledons (Fig. 2E) and primary roots (Fig. 2F) while the nYFP signal was increased in the *tni* columella cells, which appeared to have bigger nuclei (Fig. 2F).Quantitative analysis of DR5 activity by Western blot analysis using anti-GFP antibody showed much less DR5::nYFP signal in the *tni* seedlings compared to the wild-type (Col-0) (Fig. 2G). Similarly, the β-glucuronidase activity was reduced to nearly half in the*DR5::GUStni*seedlings compared to *DR5::GUS* seedlings (Fig. 2H). These results suggest that auxin response is reduced in the *tni* mutant.

**Figure 2.**
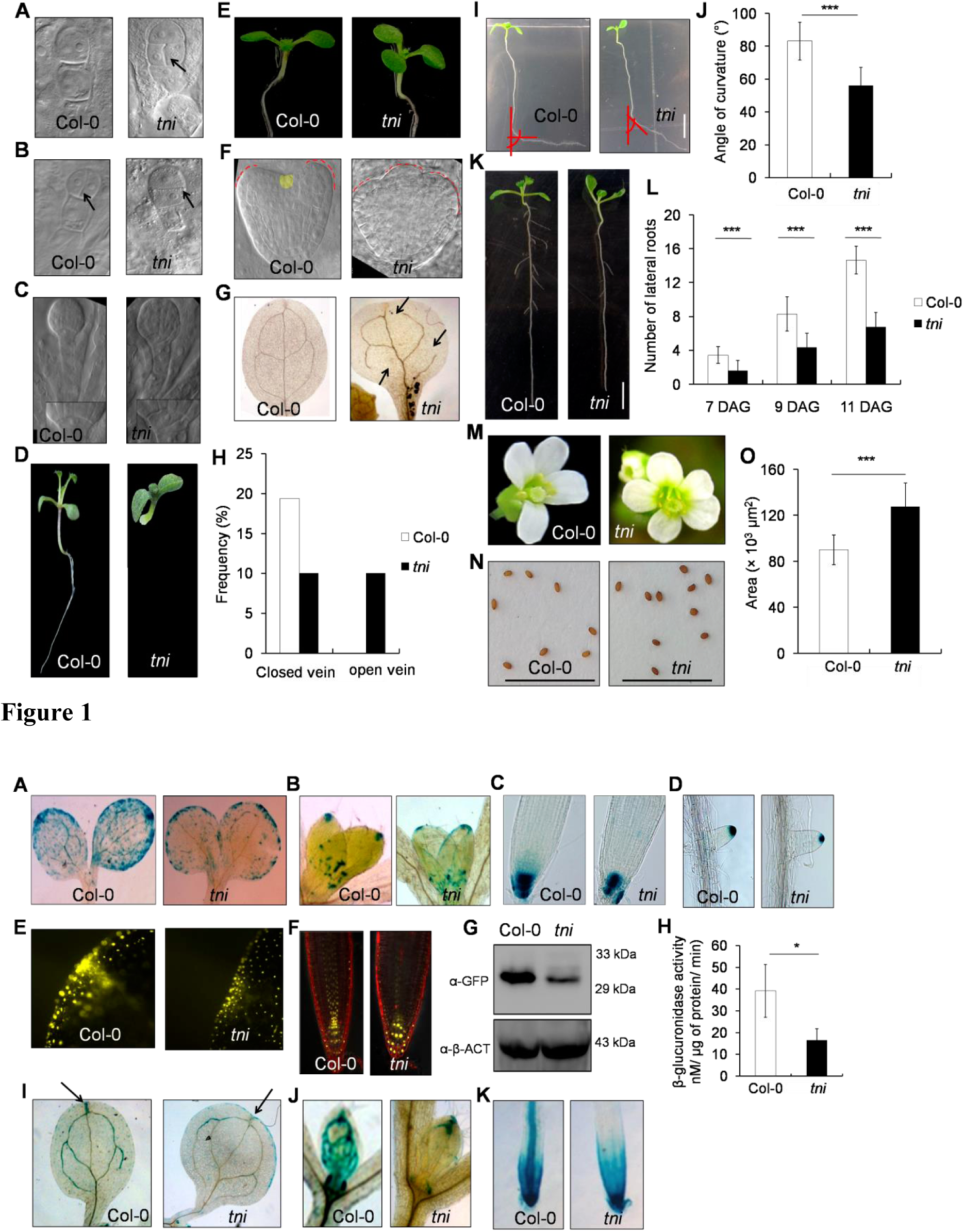
Auxin response in *tni*. **(A-D)** *DR5::GUS*activity in the cotyledons (**A**), first leaf pairs **(B)**, primary root tips **(C)** and lateral root tips **(D)** of 3-day old**(A)** and 7-8-day old (**B**-**D**) *DR5::GUS*(Col-0) and *DR5::GUS;tni*(*tni*) seedlings. **(E and F)** *DR5::nYFP*signal at the cotyledon tip of 5-day oldseedlings (**E**) and primary root tips of 7-8-day old *DR5::nYFP* (Col-0) and *DR5::nYF;tni*(*tni*) seedlings. **(F)**. Root samples were stained with propidium iodide in (**F**). **(G)** Anα-GFP Western blot analysis of the total protein extracted from 7-day old*DR5::nYFP* (Col-0) and *DR5::nYFP;tni* (*tni*) seedlings. α-β-ACTIN (α-β-ACT) was used as a loading control. Molecular weights of the protein marker were indicated on the right. **(H)**β-glucuronidase activity estimated in total extracts of*DR5::GUS* (Col-0) and *DR5::nYFP;tni*(*tni*) seedlings. Averages of biological triplicates are shown. Error bars represent SD. Statistical analysis was done by unpaired Student’s *t*-test. * indicates p=0.0406.**(I-K)***IAA2::GUS*activity in the cotyledons (**I**), first leaf pairs**(J)** and primary roots **(K)** of 7-8-day old *IAA2:GUS* (Col-0) and *IAA2::GUS;tni*(*tni*) seedlings. Arrows indicated the *IAA2:GUS* activity at the tip of the cotyledons in **(I)**.

*IAA2* is an immediate auxin-responsive gene whose induction depends on the threshold auxin level (Abel et al., 1995; Marchant et al., 2002). The venation patterning defect in *tni*, along with its reduced auxin response, prompted us to compare *IAA2::GUS* activity in Col-0 and *tni* seedlings.GUS signal was detected primarily in the vasculature of wild type cotyledons, leaves and primary roots, and in the root meristematic region (Fig. 2I-K), an expression pattern that is consistent with previous reports (Marchant et al, 2002). By contrast, the vascular *IAA2::GUS* activity was nearly absent from all these *tni*organs, with expression limited to only their tips. In addition, an ectopic expression was observed on the cotyledon margin of *tni* which was absent from Col-0 (Fig. 2I). The histochemical analysis of *DR5::GUS*and *IAA2::GUS*, together with the *DR5::nYFP* expression data, suggests that auxin response is reduced in *tni* mutant, implying a possible involvement of TNIin promoting auxin response.

The dataset of an earlier microarray experiment,carried out on the young leaves of Col-0 and *tni* (Karidas et al., 2015), was analyzed for the auxin-related genes. In keeping with the auxin related phenotypes and reduced auxin response in *tni* mutant,29 auxin-related genes were found to be differentially expressed by>2 fold (16 down-regulated and 13 up-regulated)in young *tni* leaves compared to wild type (Supplemental Fig. S2A). These up/ down-regulated genes are involved in auxin biosynthesis, transport or signaling. Many of these transcripts are also altered in seedlings externally treated with indole-3-acetic acid (IAA) (Supplemental Fig. S2B). Taken together, it appears that TNI is required to maintain normal auxin response in Arabidopsis.

### Altered sensitivity of *tni*to external auxin manipulation

The altered auxin response in *tni* could be due to perturbed auxin level or signaling. To test this, we compared the sensitivity of Col-0 and *tni* seedlings towards exogenous administration of the synthetic auxin, 1-naphthaleneacetic acid (NAA). Since auxin is known to stimulate lateral root formation in a dose-dependent manner (Ruegger et al., 1998; Ivanchenko et al., 2010),we used the number of lateral roots as a read-out of auxin sensitivity.

In Col-0, the number of lateral roots progressively increased with the increase of NAA up to 100 nM, followed by a decrease with a further increase in NAA concentration (Fig. 3A and B), thus forming a characteristic bell-shaped auxin-response curve (Ivanchenko et al., 2010). Though a similar trend was observed for *tni* roots, the peak response in *tni* was achieved at a NAA concentration (200 nM) that is twice that required for Col-0 (Fig. 3B). Even at 400 nM NAA concentration, where the relative increase in Col-0 lateral root number was only twice that of untreated plants, the *tni*lateral root number was 8-times more compared to the untreated control (Fig. 3B). Thus, the *tni* plants showed auxin response similar to wild-type but at higher hormone concentration, which is consistent with the reduced auxin response as detected by the auxin-responsive reporter assay (Fig. 2). Conversely, *tni* mutant was found to be more sensitive to the polar auxin transport inhibitor N-1-naphthylphthalamic acid (NPA) which blocks lateral root initiation by reducing the IAA level at the basal root meristem (Casimiro et al., 2001). The total number of lateral roots in Col-0 remained unaltered till 400 nM NPA, beyond which the value declined to ∼20% at 1000 nM concentration (Fig. 3C and D). In *tni*seedling, the lateral root number reduced to <40% at 400 nM NPA and to nearly zero at 1000 nM concentration. Taken together, these results suggest a reduced endogenous auxin response in *tni* roots.

**Figure 3.** Auxin sensitivity of *tni* mutant. (**A and B**) Average number of lateral rootsof 7-day old seedlings grown in the presence of 1-naphthaleneacetic acid (NAA) (**A**) and the relative increase in lateral root number upon NAA treatment (**B**). Error bars represent SD. Statistical analysis was done using unpaired Student’s *t*-test. *** denotes p≤0.0001. ns, not significant. N=12-15.**(C and D)** Average number of lateral roots in 9-dayold seedlings treated with N-1-naphthylphthalamic acid (NPA) and the relative inhibition in lateral root number upon NPA treatment (**D**). Error bars represent SD. Unpaired Student’s *t*-test was performed for statistical analysis. *** and * denote p≤0.0001 and <0.006, respectively. N=12-15.

### Genetic interaction between *tni* and mutants with auxin-related growth defects

Auxin response depends on a combinatorial effect of auxin biosynthesis, transport and signaling (Kim et al., 2007; Swarup et al., 2008; Ruegger et al., 1998; Gray et al., 1999). The molecular and genetic pathways involved in auxin response are well studied in Arabidopsis which allowed us to assess the genetic link between *TNI* and auxin pathway. We crossed *tni* with mutants defective in auxin signaling and transport such as *arf7-1*, *tir1-1*, *aux1-7* and *pin1-5* to study the phenotype in the double homozygous lines. The number of lateral root was used as a read-out to interpret the genetic interaction studies (Ruegger et al, 1998).

*ARF7* and *ARF19* redundantly regulate lateral root formation by directly activating *LBD16* (*LATERAL ORGAN BOUNDARIES-DOMAIN 16*) and *LBD29* (Okushima et al., 2007). The *arf7-1* mutant displays auxin-related phenotypes including fewer lateral roots and agravitropic response in hypocotyl (Okushima et al., 2005). Since the *arf7-1 arf19-1* double mutant totally lacks lateral root and *arf19-1*lateral root phenotype is very weak (Okushimaet al., 2005), we studied the genetic interaction of *tni* with *arf7-1*, which shows fewer lateral roots than wild-type. Lateral root formation was severely reduced in the *arf7-1 tni* double mutant compared to the individual parental lines (Fig. 4A), suggesting that *TNI* function is linked to the auxin pathway in promoting lateral root number.

**Figure 4.**
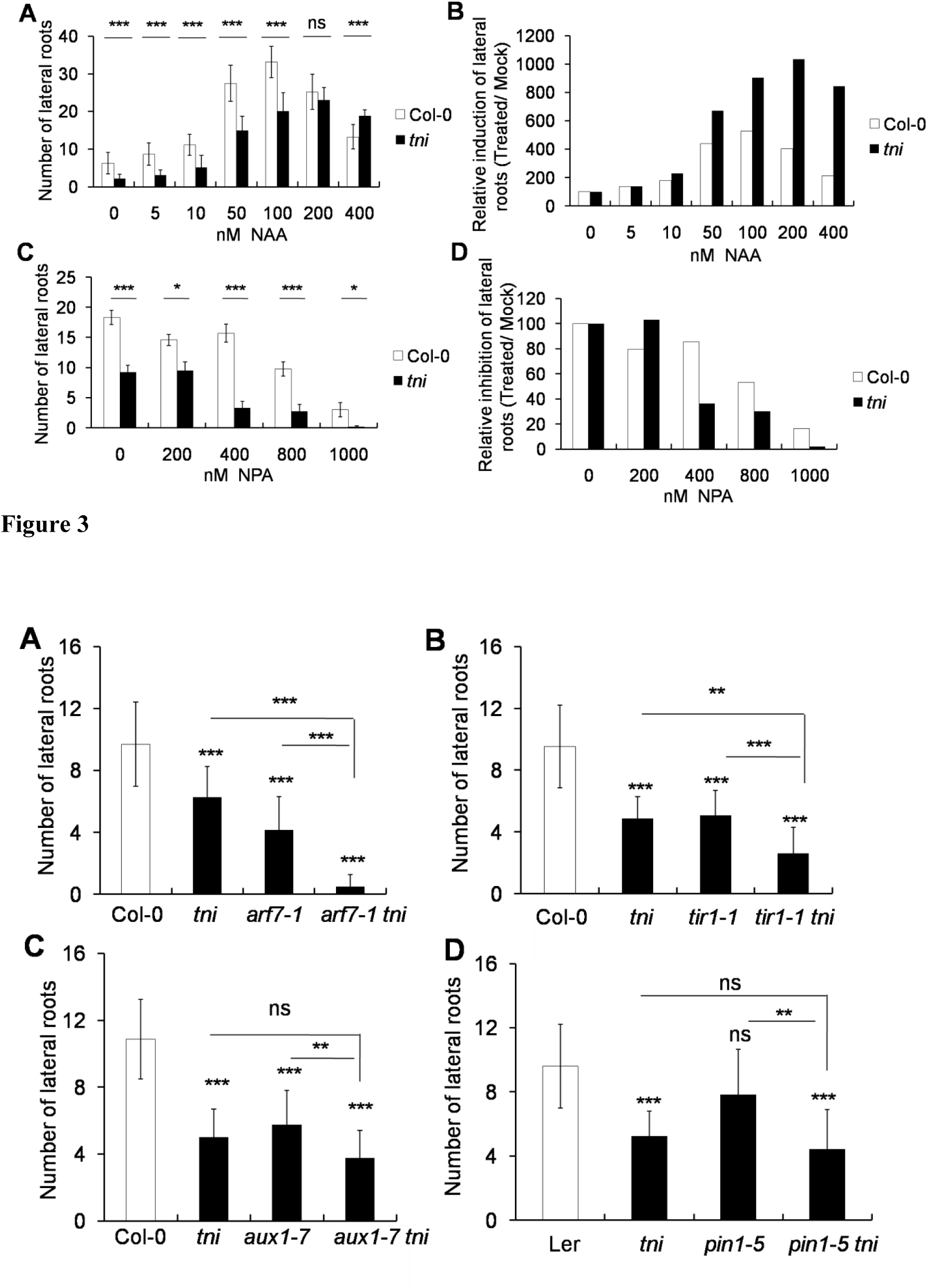
Genetic interaction of *tni* with mutants having auxin-related defects. **(A-D)** Average number of lateral roots of 9-day old seedlings. Error bars represent standard deviation (SD). The statistical analysis is done by unpaired Student’s *t*-test. *** in**(A)-(D)** denotes p<0.0001. ** denotes p= 0.0084 in **(B)** and ** denotes p= 0.0023 in **(D).**ns, not significant.N=10-15.

*TIR1* is another genetic factor that is required for normal auxin response in Arabidopsis as the *tir1-1* mutant exhibits several auxin-related growth defects such as fewer lateral roots, reduced hypocotyl elongation and moderately reduced apical dominance (Ruegger et al., 1998). The lateral root formation was reduced in *tir1-1 tni* double mutant compared to the individual parents, suggesting that these two molecules function in the same pathway in regulating lateral root formation (Fig. 4B).*AUX1* encodes an auxin influx carrier that promotes lateral root formation by facilitating the distribution of auxin from leaf to root. The *aux1-7*seedlings make fewer lateral roots and lateral root primordia (Marchant et al., 2002). Genetic interaction showed that the *tni* mutation further reduced the *aux1-7* lateral root number (Fig. 4C). In addition, genetic interaction of *tni* with the weaker allele of *pin1-5*showed that*TNI*is epistatic to *PIN1* as the *pin1-5tni*double mutant produced similar number of lateral rootsas *tni*(Fig. 4D). Together, these genetic interaction studies are in accordance with the role of *TNI* in regulating auxin response in Arabidopsis.

### *TNI* codes for UBIQUITIN SPECIFIC PROTEASE14 (UBP14)

In order to identify the *tni* locus, a map-based cloning approach was undertaken. The mutation was first trapped within a 65kb region with the help of 927 recombinant mutant plants in a mapping population (see Methods section). Sequencing of the candidate genes within this interval revealed a G→A transition in the *At3G20630* locus (Fig. 5A and Supplemental Fig. 3A and B). This mutation mapped at the canonical 3′ splice acceptor site at the junction of the 3^rd^ intron and the 4^th^ exon, which is necessary for intron recognition by the spliceosome (Sharp et al., 1987). *At3G20630*is predicted to encode UBIQUITIN-SPECIFIC PROTEASE 14, a ZnF de-ubiquitinase protein involved in ubiquitin recycling(Doelling et al., 2001; Tzafrir et al., 2002; Xu et al., 2016). Several alleles of *At3G20630* had been previously described, most of which show embryonic lethality (Doelling et al., 2001; Tzafrir et al., 2002) (Fig. 5A). To further test the identity of the *tni* locus, we performed allelism test with *titan6-4* (*ttn6-4*) plants, one of the known alleles of *UBP14*(Tzafrir et al., 2002). The *tni* homozygous plants were crossed to *ttn6-4* heterozygous plants(which resembled Col-0) and plants with cup-shaped rosette leaves were observed in the F_1_ generation (Fig. 5B-D),while the *tni* plants crossed to Col-0 produced all wild-type looking individuals (data not shown), suggesting that *tni*is allelic to *ttn6*. Further, the cup-shaped phenotype of *tni*leaves was rescued by over-expressing the wild-type *TNI*transcript in the *tni*background(3 out of 14 hygromycin-resistant transformants recovered, produced completely flat rosette leaves) (Fig. 5E). In addition, reducing the *AtUBP14*transcript in transgenic lines expressing an artificial micro RNA against the wild-type *TNI* transcript under the constitutive *RPS5a* promoter recapitulated *tni* phenotype (Fig. 5F). Three out of 25 hygromycin-resistant transformants recovered produced rosette leaves with weak cup-shaped phenotype.

**Figure 5.**
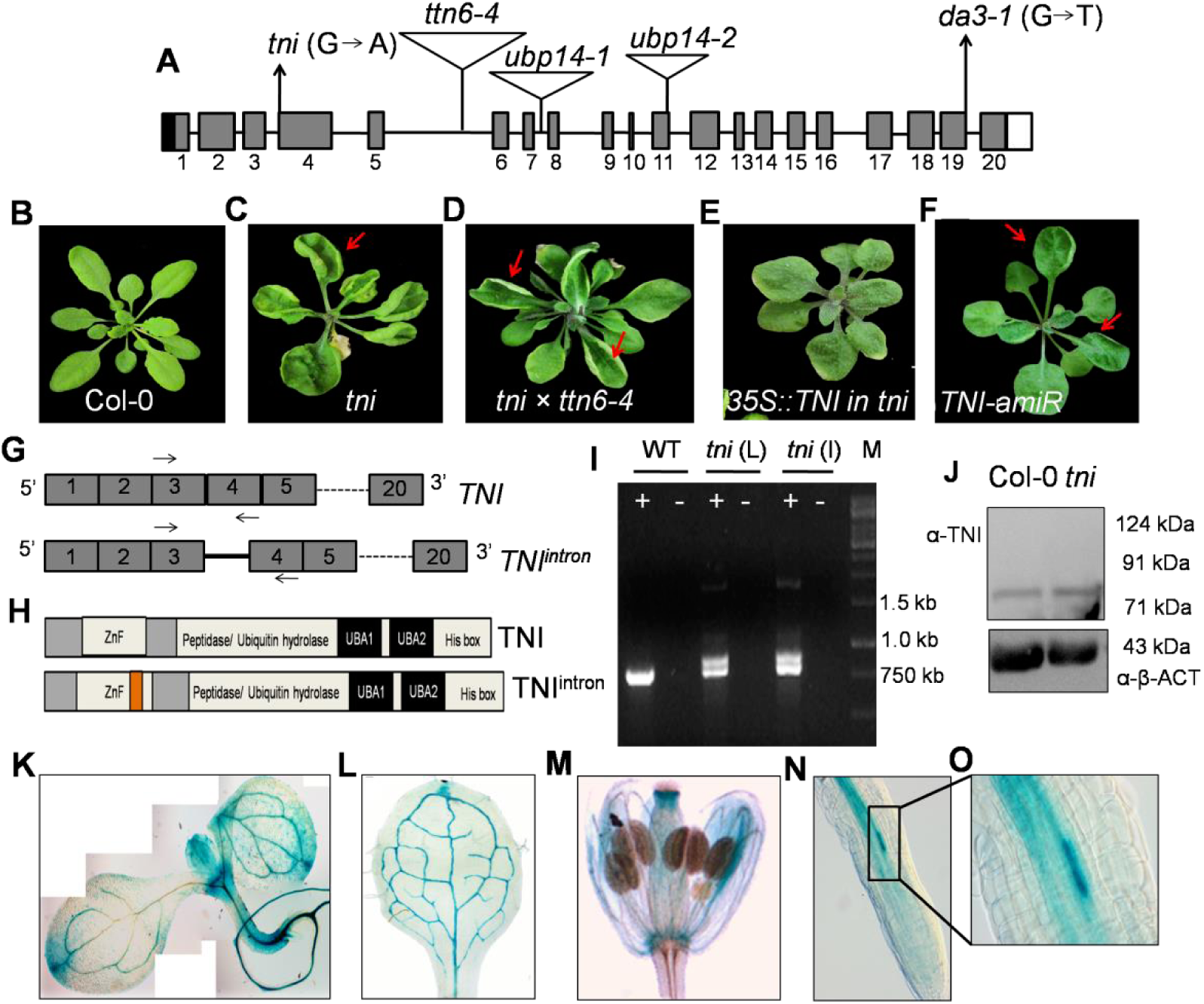
Cloning of *TNI*. **(A)** A schematic representation of the *TNI*/ *UBP14*genomic locus showing the 5′ UTR (black box), twenty exons (grey box), 19 introns (black line) and the 3′ UTR (white box). Positions of the T-DNA insertion in various alleles of *UBP14* are shown by open inverted triangles. The *G A* transition at the third intron-exon junction in the *tni* allele and the *G T*substitution at the nineteenth exon-intron junction in the *da3-1* allele are indicated. **(B)** 30-dayold rosettes of Col-0, *tni* **(C)** and *tni* x *ttn6-4* (+/-) F1 plant **(D)** to show that *TNI* and *TTN6* form a single complementation group. Red arrows indicate the curvature in *tni* leaf and *tni* x *ttn6-4* in F1 genaration.**(E)** Rosette of 30-day old *tni* plant over-expressing *TNI* shows rescue in *tni*phenotype.**(F)**A 30-day old Col-0 plant expressing an artificial microRNA (amiR) targeting the *TNI* transcript produced cup shaped leaf as indicated by red arrows.**(G)** Schematic representation of the predicted wild-type *TNI* transcript with all the introns efficiently spliced out (upper panel) and the mutant transcript (*TNI^intron^*) showing retention of the 3^rd^ intron (black line). Arrows indicate the position of the primers on the genomic locus used for RT-PCR analysis.The broken lines indicate the continuity up to the 20^th^ exon in the transcripts. **(H)**The domain architecture of TNI protein with N-terminal ZnF domain, peptidase domain along with C-terminal UBA domains (upper panel) and TNI^intron^ showing insertion of 34 amino acids in the ZnF domain indicated by orange box. **(I)**An ethidium bromide-stained agarose gel showing the products of RT-PCR using the total RNA from Col-0 (WT), mature *tni* leaf (*tni* L) and *tni* inflorescence (*tni* I). +and - indicate cDNA and RNA as PCR template. M indicates 1 kb DNA marker. **(J)** Western blot analysis of the total protein extracted from 7-day old seedlings showing the bands corresponding to endogenous TNI protein of predicted molecular weight 88 kDa using anti-TNI polyclonal antibody. Anti-β-ACT was used as loading control. **(K)**A 8-day old transgenic seedling expressing *pTNI::GUS*.**(L)**The 1^st^ leaf of11-day old *pTNI::GUS* seedling shows the *TNI* promoter activity in the vasculature. (M) *pTNI::GUS* expression in the floral organs.**(N)**Primary root of 8-day old *pTNI::GUS* seedling shows GUS activity in the vasculature. The inset in **(O)** shows the expression of *TNI* promoter in the pericycle cell magnified in **(N).**

The 5340 nucleotide long *TNI*primary transcript consists of 20 exons and 19 introns that yield a 2394 nucleotides long mature transcript predicted to encodea 88 kDa protein product (Fig. 5A; Supplemental Fig.S3C). If the G→A mutation in the *tni*locus interferes in the splicing process, it is predicted to lead to an in-frame insertion of the 102 nucleotide long 3^rd^ intron in the mature transcript, resulting in an aberrant mature transcript (*TNI^intron^*) which would encode a full-length TNI protein with an extra 34 residue insertion at the 127^th^ residue downstream to the N-terminus of TNI protein (Fig. 5G and H; Supplemental Fig. 3D). While RT-PCR analysis with primers flanking the 3^rd^ intron (Fig. 5G) detected a single product of 750 bps in both Col-0 and *tni*leaves, an additional product corresponding to the retention of the 3^rd^ intron was detected in the *tni* leaves (Fig. 5I), implying that this intron is retained in *TNI^intron^*. Comparable intensity of the two bands corresponding to *TNI* and *TNI^intron^* in the RT-PCR analysis indicates nearly equal abundance of the two transcripts in the *tni* mutant plants. The endogenous TNI protein was detected in both Col-0 and *tni* by Western blot analysis using an anti-TNI antibody generated against a TNI-specific peptide present in both wild-type and mutant proteins (Fig. 5J; Supplemental Fig. 3E), further suggesting that *tni*is not a null allele. The anti-TNI antibody could not distinguish TNI^intron^from TNI in the *tni*sample of the Western blot since the mutant protein is predicted to be merely 4% larger than the wild type protein.

As the pleotropic *tni*phenotype was observed throughout the mutant individual, we examined the *TNI* promoter activity in transgenic plants expressing a*pTNI::GUS* reporter cassette. In all the five independent *pTNI::GUS*transgenic lines obtained, and GUS activity was detected almost throughout the seedlings with intense signal observed in young leaves, cotyledon tips, shoot apex, primary roots as well as in floral organs (Fig. 5K-O). Signal was restricted to the vasculature as the leaves matured (Fig. 5L). GUS activity was detected in petals and carpels as well (Fig. 5M). In primary roots, the GUS signal was restricted mostly in the vasculature with more intense signal in discrete pericycle cells that are known to initiate lateral roots (Fig. 5N and O) (De Smet et al., 2012). This expression pattern is in general agreement with the expression of UBP14 reported earlier using western blot analysis or GUS reporter assay (Doelling et al., 2001; Xu et al., 2016).

### Cell type-specific TNI activity regulates lateral root formation

The root phenotype observed in *tni* such as reduced lateral root number (Fig. 1K and L) could be due to the local promoter activity or due to the systemic effect of altered auxin response. To test this, we manipulated *TNI*activity in the lateral root initials, since *TNI* promoter is highly active in specific pericyle cell-types (Fig. 5N and O), and studied its effect on lateral root growth.To test the level of wild-type *TNI* on lateral root growth, *TNI*transcript was down-regulated specifically in the lateral root primordium using an artificial microRNA (amiR) against the endogenous *TNI* transcript (Supplemental Fig. S4A). A truncated *PLETHORA7* (*PLT7*) promoter (*pPLT7*) which is specifically active in the lateral root primordia was used for the study (Prasad et al., 2011).The number of lateral root was 45-80% reduced in six independent *pPLT7::TNI-amiR* transgenic lines in the T_2_ generation (data not shown).Reduced lateral root number was also observed in the two homozygous *pPLT7::TNI-amiR* transgenic lines(Fig. S4B and C) established in the T_3_ generation. These results suggest that the reduced lateral root phenotype of the *tni* mutant is due the local loss of TNI activity in the lateral root initials and not due to a systemic effect of altered auxin response.

When the TNI^intron^ protein was expressed in the lateral root initials (Supplemental Fig. S4A), the number of lateral roots was reduced to ∼50% in a homozygous *pPLT7::TNI^intron^* transgenic line, a reduction similar to what was observed in the *tni* allele (Fig. S4B and D). This suggests that the reduced lateral root in*tni* is due to the local expression of the aberrant *TNI^intron^* transcript.

### Increased accumulation of poly-ubiquitin and poly-ubiquitinated proteins in *tni*

In Arabidopsis, UBP14 consists of an N-terminal ZnF domain, a peptidase domain and two C-terminal consensus ubiquitin-associated UBA domains (Fig. 5H) (Yan et al., 2000; Doelling et al., 2001). UBP14 has been reported to hydrolyze both α-linked and iso-linked poly-ubiquitin chains into mono-ubiquitin (Doelling et al, 2001; Xuet al., 2016). Since *TNI* encodes UBP14, we compared the ubiquitinated protein profile of Col-0 and *tni*. Western blot analysis of the total protein samples from Col-0 and *tni*plants using anti-ubiquitin antibody showed an excess accumulation of higher molecular weight poly-ubiquitinated proteins and reduced amount of mono-ubiquitin in *tni* compared to Col-0 (Fig. 6A; Supplemental Fig. S5A). In addition, free poly-ubiquitin chains corresponding to tri-and tetra-ubiquitin were detected in *tni* while the Col-0 sample showed no poly-ubiquitin accumulation (Fig. 6A), implying that TNI is involved in ubiquitin recycling in Arabidopsis which is perturbed in the *tni* mutant. Poly-ubiquitin chains of distinct linkages are identified in Arabidopsis, of which the most abundant Lys48-linked poly-ubiquitin is implicated in protein turn-over by 26S proteasome while Lys63-linked poly-ubiquitin imparts non-degradative fate to the cellular proteins (Kim et al., 2013; Mevissen and Komander, 2017). To determine the precise role of TNI in cellular ubiquitin homeostasis, we examined the specific ubiquitin linkage in the poly-ubiquitinated proteins that accumulated in the *tni* mutant. Western blot analysis of the total protein extracts from Col-0 and *tni* seedlings using ubiquitin linkage-specific antibodies showed a higher abundance of Lys48-linked poly-ubiquitinated proteins in *tni* relative to Col-0 (Fig. 6B) while that of Lys63-linked poly-ubiquitinated proteins appeared to be similar to that of Col-0 (Fig. 6C). This suggests that TNI is involved in the turn-over of the cellular proteins by Ubiquitin-26S proteasomal pathway.

**Figure 6.**
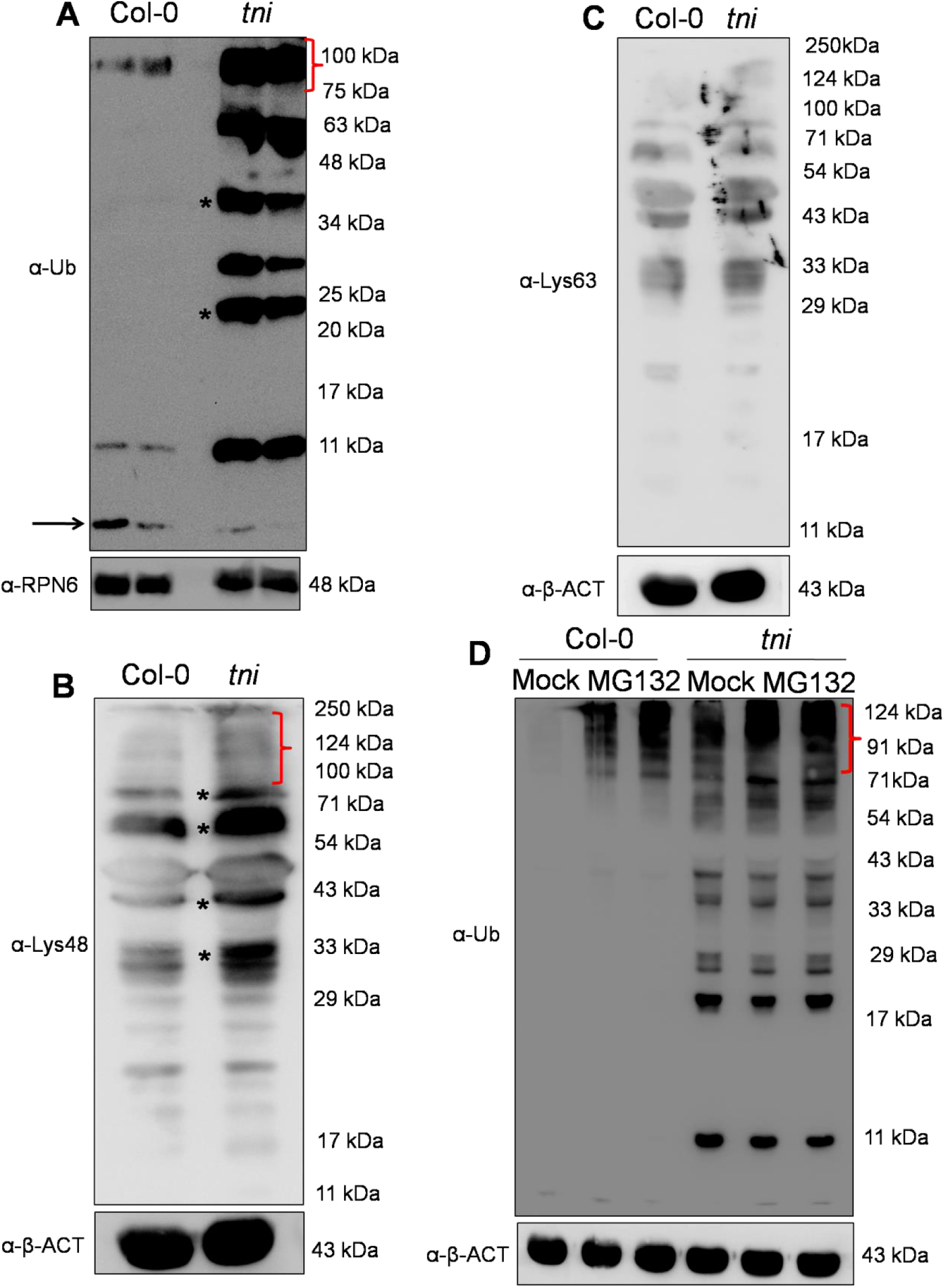
Detection of poly-ubiquitinated proteins in *tni* mutant. **(A)** An anti-ubiquitin (-Ub) Western blot analysis of total proteins extracted from 7-day old seedlings. Ubiquitin polymers are indicated by *. The red bracket indicates the accumulation of higher molecular weight poly-ubiquitinated proteins. The level of free mono-ubiquitin is marked by the black arrow.The -RPN6 Western blot analysisserved as loading control. Numbers indicate molecular weight of the protein marker. **(B)**A Western blot of total proteins extracted from 7-day old seedlings probed with α-Lys48 ubiquitin antibody. Smears within the red bracket mark the ubiquitinated targets proteins of higher molecular weight. Asterisks denote accumulation of Lys48 linked poly-ubiquitinated target proteins. Anti-β-ACT Western blotanalysis served as a loading control. Numbers indicate molecular weight of the protein marker. **(C)**An α-Lys63 ubiquitin Western blot analysis of total proteins extracted from 7-day old seedlings. The α-β-ACT Western blot served as loading control. Numbers in the right indicate molecular weight of the protein marker. **(D)** An -Ub Western blot of total proteins extracted from 7-day old seedlings and treated with 0 (Mock) (1^st^ and 4th lanes), 0.1 (2^nd^ and 5^th^ lanes) or 0.2 mM (3^rd^ and 6^th^ lanes) MG132 for 16 hrs. Smears marked by a red bracket indicate poly-ubiquitinated target proteins. Anti-β-ACT Western blot served as a loading control. Molecular weight of the protein markers are shown on the right. The protein samples were resolved in 15% SDS-PAGE in **(A - D)**.

The inefficient turn-over of poly-ubiquitin chains and the accumulation of poly-ubiquitinated proteins is likely to render the *tni* mutant more sensitive to MG132,a peptide aldehyde that acts as a reversible inhibitor of 26S proteasome by blocking the catalytic residues in the β-subunits of the protease (Kisselev and Goldberg, 2001). To test this, we comparedMG132 sensitivity of Col-0 and *tni*by detecting the accumulation of poly-ubiquitinated proteins in the presence of this inhibitor. The mock-treated *tni* sample showed more ubiquitinated protein signal than mock-treated Col-0, and the intensity of the signal in *tni*increased to a level higher than the Col-0 value at all concentrations of MG132 used (Fig. 6D; Supplemental Fig. S5B). This result suggests that the *tni* cells accumulate more ubiquitinated target proteins and are hyper-sensitive to a perturbation of proteasome activity. MG132 treatment didnot result in an accumulation of polyubiquitin chains in Col-0 neither did it alter the steady-state level of poly-ubiquitins in the *tni* sample (Fig. 6D). Taken together, these observations suggest that the accumulation of free poly-ubiquitin chains in *tni*is likely due to an inefficient hydrolysis of its substrates, leading to a sub-optimal level of functional TNI protein insufficient to disassemble the unanchored poly-ubiqutin chains to mono-ubiquitin.

### Lack of de-ubiquitination activity of TNI^intron^ towards poly-ubiquitin chains

The *tni* mutant is predicted to contain both TNI and TNI^intron^forms of the protein. Thus, the accumulation of poly-ubiquitins and poly-ubiquitinated target proteins could be due to an inadequate catalytic activity of TNI, which are otherwise processed efficiently by the ubiquitin proteases (Doelling et al., 2001). To test this, we compared the activity of wild-type TNI and mutant TNI^intron^ proteins towards their substrate by *invitro* de-ubiquitination assay (Doelling et al., 2001; Rao-Naik et al., 2000). Recombinant TNI efficiently cleaved 2-7-mer Lys48-linked poly-ubiquitin substrates into mono-ubiquitin whereas TNI^intron^failed to produce any mono-ubiquitin product (Fig. 7A). As a positive control, we generated a TNI^C317S^ form of the protein which contains a Cys→Ser point mutation at the active site residue C317. This point mutation has been reported to render *At*UBP14 catalytically inactive (Doelling et al., 2001). Like TNI^intron^, TNI^C317S^also failed to cleave the poly-ubiquitin substrates. De-ubiquitination assay was also performed *invivo* within *E. coli* cells that co-expressed α-linked hexa-ubiquitin chain encoded by a *UBQ10*construct (Rao-Naik et al., 2000) in the presence of various forms of the TNI proteins. While hexa-ubiquitin was completely cleaved into di- and mono-ubiquitin forms by wild-type TNI and *Sc*UBP14, a functional homolog of TNI in yeast (Amerik et al., 1997), TNI^intron^ and TNI^C317S^failed to cleave the *UBQ10*-encoded hexa-ubiquitin (Fig. 7B). A similar *invivo* de-ubiquitination assay in *E.coli*expressing α-linked, His-tagged tetra-ubiquitin (Rao-Naik et al., 2000) also showed that TNI, but not TNI^intron^ and TNI^C317S^, cleaves the substrate into His-tagged di- and mono-ubiquitin products as detected by anti-His antibody (Supplemental Fig. S5C). Thus, TNI^intron^is catalytically inactive towards poly-ubiquitin substrates.Therefore, it is possible that the accumulation of poly-ubiquitinated proteins of higher molecular weight in *tni*is a consequence of improper disassembly of free poly-ubiquitin chains. Incubation of total protein extract from *tni* seedlings with recombinant TNI resulted in the disappearance of the free poly-ubiquitin chains (Fig. 7C), converting the Western blotprofile somewhat similar to the Col-0 profile shown earlier (Fig. 6A). However, recombinant TNI^intron^ and TNI^C317S^ did not hydrolyze these multi-ubiquitin chains, further demonstrating their inability to hydrolyze these substrates.

**Figure 7.** TNI^intron^ is catalytically inactive. **(A)** An anti-ubiquitin (α-Ub) Western blot of Lys48-linked 2-7 unit poly-ubiquitin chains (input) incubated with purified recombinant GST-TNI (TNI), GST-TNI^intron^(TNI^intron^) and GST-TNI^C317S^(GST-TNI^C317S^) fusion proteins. Arrow indicates mono-ubiquitin band. Free GST served as a negative control.The α-GST Western blot (lower panel) serves as loading control. Numbers indicate molecular weight of the protein marker. **(B)**An α-UbWestern blot of *E. coli*lysate expressing *UBQ10* encoding hexa-ubiquitin (Ub_6_) along with recombinant GST-*Sc*UBQ14 (*Sc*UBQ14), GST-TNI (TNI), GST-TNI^intron^(TNI^intron^) and GST-TNI^C317S^(TNI^C317S^) fusion proteins. Purified mono-ubiquitin (Mono Ub) was loaded as a positive control. Lysates expressing no recombinant proteins (BL21), recombinant GST alone (GST) and recombinant GST plus UBQ10 were used as negative controls. Numbers on the right indicate molecular weights of marker proteins. A Ponceau-stained membrane shown below served as loading control. **(C)** An α-Ub Western blot analysis of total protein extracted from 7-dayold *tni* seedlings treated with various forms of TNI protein. The input in lane 1 corresponds to the *tni* protein extract which shows several poly-ubiquitin bands that are cleaved by the wild-type TNI protein (lane 3) but not by GST alone (lane 2), TNI^intron^ (lane 4) or by GST-TNI^C317S^ (lane 5).

### *In vitro* substrate binding of TNI^intron^

It is possible that the 34-residue insertion in the ZnF domain of TNI^intron^ interferes either with its substrate-binding ability or with its catalytic activity *per se*(Fig. 5H). To test this, we compared the substrate binding property of the full-length TNI and its mutated versions, TNI^intron^ and TNI^C317S^, with Lys48-linked tetra-ubiquitin (Fig. 8A). A minimum tetra-ubiquitin chain linked with Lys48 linkage is sufficient forefficient targeting of proteins for degradation by 26S proteasome (Thrower et al., 2000) and our data suggests that TNI works in the 26S proteasomal pathway. With this rationale, we used only tetra-ubiquitin as a substrate for binding assay. We observed that both wild-type and mutant forms of the protein – TNI, TNI^intron^ and TNI^C317S^bound to the substrate, while only the wild-type form cleaved Lys48-linked tetra ubiquitin into mono-ubiquitin (Fig. 8A). Thus, retention of the 3^rd^ intron in TNI^intron^ does not interfere with its substrate binding property.

**Figure 8.**
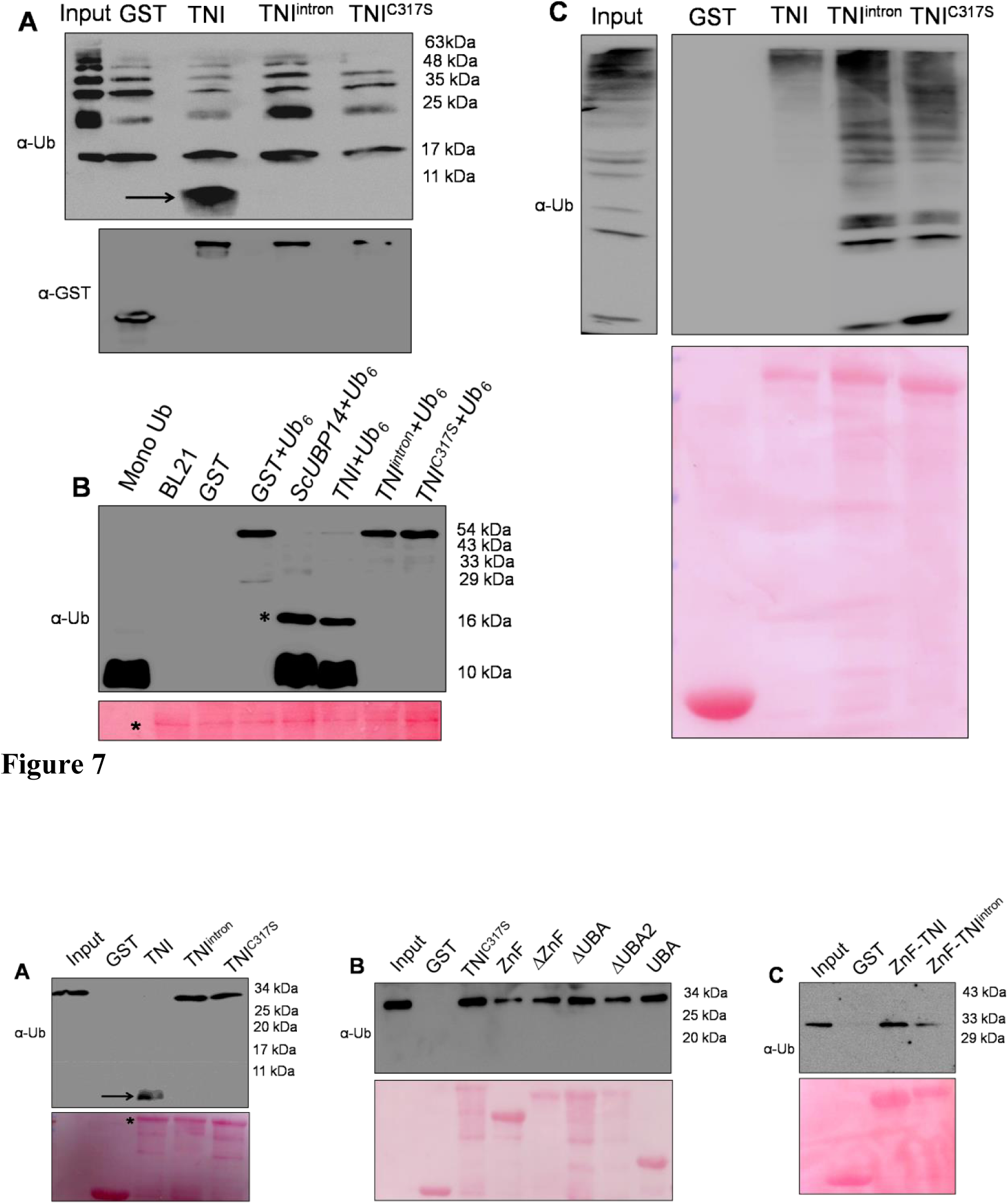
Ubiquitin binding by various forms of TNI protein. **(A)** An anti-ubiquitin (α-Ub) Western blot of Lys48-linked tetra-ubiquitin treated with recombinant GST-TNI (TNI), GST-TNI^intron^ (TNI^intron^) and GST-TNI^C317S^ (TNI^C317S^) fusion proteins expressed in *E.coli* cells. GST, GST-TNI, GST-TNI^intron^ and GST-TNI^C317S^ were immobilized by glutathione beads and incubated with purified Lys48-linked tetra-ubiquitin in binding buffer.Lys48-linked tetra-ubiquitin alone (Input) and only GST protein served as a positive and negative controls respectively. TNI bound and cleaved the substrate into mono-ubiquitin as indicated by black arrow. TNI^intron^ and TNI^C317S^ could bind the substrate efficiently (compared to input) but did not hydrolyze the substrate. Numbers on the right indicate molecular weights of the protein marker. A Ponceau-stained membrane shown below served as loading control. **(B)**An α-Ub Western blot analysis of the *in vitro* substrate binding assay with indicated GST-tagged deletion forms of TNI (labels above the lanes) and Lys48-linked tetra-ubiquitin. The blot shows that all the deletion versions of TNI bind to tetra-ubiquitin substrate. GST serves as a negative control (lane 2). Input in lane 1 shows Lys48-linked tetra-ubquitin used as a substrate. A Ponceau-stained membrane served as loading control. **(C)** An α-ubiquitin Western blot analysis of *in vitro* substrate binding assay shows that GST-tagged version of ZnF-TNI and ZnF-TNI^intron^ bind to tetra-ubiquitin while GST alone (negative control) does not bind. A Ponceau-stained membraneserved as loading control.

It had been reported earlier that the two tandem UBA domains in UBP14 are essential but not sufficient for ubiquitin binding (Xu et al., 2016). However, the exact UBP14 domains involved in binding poly-ubiquitin are not clear. To map the poly-ubiquitin binding domains of TNI, we generated and expressed in *E. coli* the following five truncated forms of TNI (Supplemental Fig. S6A-C): (i) the N-terminal ZnF domain alone (ZnF), (ii) TNI without theZnF domain (ΔZnF), (iii) TNI without the C-terminal UBA domains (ΔUBA), (iv) TNI without the C-terminal UBA2 domain (ΔUBA2), and (v) the C-terminal tandem UBA domains alone (UBA). Since the full-length TNI efficiently hydrolyzed poly-ubiquitin into it monomer form (Fig. 8A) and thus cannot be used for comparison of ubiquitin binding by these deletion mutants, we used TNI^C317S^ (Fig. 8A) as thepositive control. In a Western blot analysis of *invitro* substrate-binding assay, all the truncated forms of TNI and TNI^C317S^bound to the Lys48-linked tetra-ubiquitin substrate with varying efficiency (Fig. 8B).It is possible that the positional context of ZnF and UBA in the folded protein, along with the peptidase domain, contribute to efficient substrate binding. Since the ZnF domain in TNI^intron^(ZnF-TNI^intron^) is disrupted by the insertion of a 34-residue fragment (Supplemental Fig. S6A), we argued that the C-terminal UBA domain might contribute to the ubiquitin binding by TNI^intron^, as the two UBA domains in isolation are capable of binding ubiquitin. Hence, we expressed the ZnF domain of TNI^intron^, ZnF-TNI^intron^and tested its substrate binding capability. In an *invitro* assay, ZnF-TNI^intron^ bound to the tetra-ubiquitin substrate (Fig. 8C), suggesting that the disrupted ZnF domain by the inclusion of the 3^rd^ intron retains its substrate-binding capability. It is possible that the inability of TNI^intron^to hydrolyze poly-ubiquitin substrate is due to an overall conformational change rendering the catalytic domain inactive.

### Stability of AUX/IAA transcriptional repressors in *tni*

The auxin-responsive AUX/IAA repressors are degraded by ubiquitin-mediated 26S proteasomal pathway and provide robustness to the auxin signaling pathway by relieving ARFs from repression (Gray et al., 2001; Leyser 2018).The increased amount of free poly-ubiquitin chains, poly-ubiquitinated proteins and the increased sensitivity of *tni* plants towards MG132 (Fig. 6A,B and D) led us to hypothesize that the *tni* mutation affects the stability of AUX/IAAs, resultingin an array of auxin-related growth defects similar to the gain-of-function mutants of AUX/IAAs with their increased stability (Tian and Reed, 1999; Hamann et al., 1999, Uehara et al., 2008). To test this, we established the *DII:VENUS* reporter line (Brunoud et al, 2012) in the *tni* mutant background and compared its DII:VENUS signal with that in wild-type.Weak VENUS signal was detected in Col-0 primary root due to a rapid turn-over of DII:VENUS (Fig. 9A, left panel), suggestive of high auxin activity (Brunoud et al, 2012). However, the Col-0 rootsexpressing a mutant, non-degradable form of the protein, mDII:VENUS showed strong and widespread VENUS signal (Fig. 9A, middle panel), which is consistent with previous reports(Brunoud et al, 2012). The *DII:VENUS tni* primary roots showed VENUS signal much stronger than Col-0 (Fig. 9A, right panel) and somewhat weaker than the *mDII:VENUS* roots. Western blot analysis of protein extract from 7-dayold seedlings using anti-GFP antibody further confirmed that the level of DII:VENUS was indeed more in the *tni* seedlings than in Col-0 (Fig. 9B). This suggests that the DII domain, and hence some of the AUX/IAA repressors, are more stabilized in *tni* roots than in Col-0. To test this, we compared the levels of IAA18 and AXR3/IAA17 proteins in Col-0 and *tni* plants using their respective translational fusion lines *IAA18:GUS* and *HS::AXR3-NT:GUS* (Ploense et al, 2009; Gray et al, 2001). The *HS::AXR3-NT:GUS* reporter line had been extensively used to monitor the turn-over of IAA17 in various mutants where protein degradation by 26S proteasome is affected (Gray et al., 2001; Stuttmann et al., 2009).Stabilization of *HS::AXR3-NT:GUS* in the *ubc13* (*ubiquitin-conjugating enzyme 13*)mutant resulted in reduced auxin response (Wen et al., 2014). *UBC13* encodes Ubiquitin-conjugating E2 enzyme which is the second enzyme in the E1-E2-E3 cascadeand is involved in protein turn-over by 26S proteasome (Wen et al., 2014; Callis, 2014). Further, the degradation of the N-terminal AXR3 (AXR3-NT) is auxin-dependent and requires 26S proteasomal activity (Gray et al., 2001). We made use of this reporter line and tested the stability of AXR3 in the *tni* mutant. GUS assay was performed in the *HS::AXR3-NT:GUS*, *HS::axr3-1-NT:GUS* and *HS::AXR3-NT:GUStni* seedlings after a 2 hr pulse of heat shock at 37°C. In Col-0 cotyledons, weak GUS signal was detected 20 min after heat shock treatment, whichcompletely disappeared within 80 min (Fig. 9C, upper panel), suggesting a rapid turnover of the protein in wild-type. As expected, the non-degradable form of the protein, axr3-1-NT:GUS accumulated in large amount at 20 min and continued to accumulate, producing strong GUS signal after 80 min (Fig. 9C, middle panel). The AXR3-NT:GUS signal in the *tni* cotyledons also accumulated in larger amount than in Col-0, and decreased at a slower pace retaining considerable signal even after 80 min of heat shock induction (Fig. 9C, lower panel). These results suggest that the IAA17/AXR3 protein is stabilized in the *tni* plants, implying a slower turn-over of this protein by 26S proteasome.

**Figure 9.**
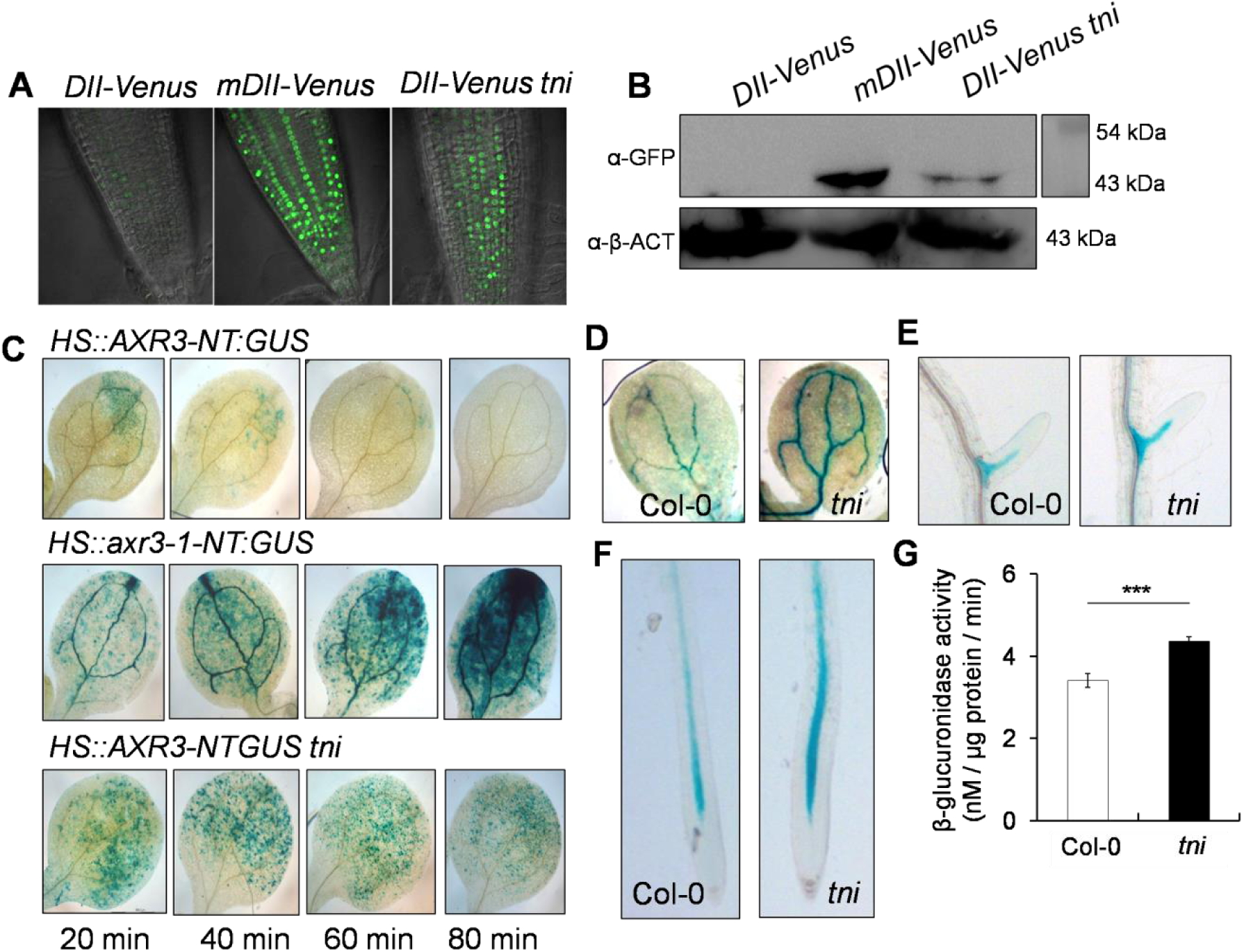
Stabilization of AUX/IAA transcriptional repressors in *tni*. **(A)** DII:VENUS signal in the primary roots of 7-8-day old Col-0 (*DII:VENUS* and *mDII:VENUS*) and *tni* (*DII:VENUStni*) seedlings. Strong signal of mDII:VENUS, a non-degradable version of DII domain served as a positive control. **(B)**An -GFP Western blot of the total protein extracts from 7-day old Col-0 (*DII:VENUS* and *mDII:VENUS*) and *tni*(*DII:VENUStni*)seedlings. Anti-β-ACT Western blottingservedas an internal control. Numbers indicate molecular weights of the protein marker. **(C)** GUS activity in the cotyledons of 7-day old seedlings of *HS::AXR3-NT:GUS* (upper panel), *HS::axr3-1-NT:GUS* (middle panel) and *HS::AXR3-NT:GUS tni* (lower panel) at various durations after heat shock treatment. The non-degradable form *axr3-1-NT* served as a positive control. NT denotes N-terminal domain. **(D)**-**(F)**IAA18:GUS activity signal in the cotyledon vein (**D**), lateral root (**E**) and primary root (**F**) of Col-0 (*IAA18:GUS*) and *tni* (*IAA18:GUS tni*) seedlings. **(G)** β-glucuronidase activity in the Col-0 (*IAA18:GUS*) and *tni* (*IAA18:GUS tni*) seedlings determined by GUS assay.Averages of biological triplicates are shown. Error bars represent SD. Statistical analysis was done by unpaired Student’s *t*-test.*** denotes p<0.0001.

The *tni* mutant resembles the gain-of-function mutants of several AUX/IAA repressors having perturbed auxin response (Hamann et al., 1999; Tian and Reed 1999; Uehara et al., 2008). One such gain-of-function mutant, *iaa18-1*hasembryo patterning defects, reduced sensitivity to exogenous auxin application and fewer lateral roots(Ploense et al, 2009; Uehara et al., 2008).To examine the accumulation of IAA18 in the *tni* mutant, we compared its levels in the wild type and the mutant seedlings. We detected IAA18:GUS activity in the vasculature of cotyledons, lateral root and primary root in Col-0 seedlings (Fig. 9D-F). Even though the pattern of the signal remained moreorless similar in the vasculature of *tni* seedlings, its intensity increased, suggesting an increased stability of IAA18 in the mutant. Quantification of β-glucuronidase activity also confirmed an increase in IAA18:GUS in *tni*plants (Fig. 9G). All these observations suggest that TNI is required for the turn-over of some AUX/IAA proteins involved in regulating auxin response.

## DISCUSSION

The *tni* allele of *UBP14*is recessive and likely hypomorphic in nature as multiple knockout lines show embryonic lethality. It exhibits diverse phenotypic defects in auxin-related processes both at the embryonic and post-embryonic stages of development. Auxin is implicated in regulating multiple aspects of plant developmental processes including early embryogenesis (ten Hove et al., 2015). The pleiotropic auxin-related phenotypes of *tni* mutant suggest that *TNI/UBP14* is likely to be a global regulator of plant growth and development. The previously reported *ttn6-4*allele of *UBP14*, a T-DNA insertion mutant, has a deletion of 400 bp resulting in the elimination of exon 6 and 7. The *ttn6-4*allele exhibits defective embryos arrested at the globular stage (Tzafrir et al., 2002). T-DNA insertion in the 7^th^ and 11^th^ intron in the *ubp14-1*and *ubp14-2*alleles respectively, results in null mutants with similar developmentally arrested phenotype at the globular stage (Doelling et al., 2001). The *tni* allele also exhibited partial embryo lethality phenotype resulting in reduced seed setting (Premananda Karidas, PhD Thesis, 2014). The aborted embryo of *ubp14-1* and *ubp14-2* had heightened levels of poly-ubiquitins and poly-ubiquitinated proteins, implying that *UBP14* is required for ubiquitin recycling which is crucial for the progression of embryo development. Our data shows that the *tni*plants have elevated accumulation of un-anchored poly-ubiquitin chains corresponding to tri- and tetra-ubiquitin as well as poly-ubiquitinated target proteins with a concomitant reduction in the free mono-ubiquitin. Thus, TNI/ UBP14 is involved in ubiquitin recycling during post-embryonic development as well, failure of which results in multiple growth and developmental defects. Detection of mono-ubiquitin in *tni*, albeit at a low abundance, implies that partial de-ubiquitination activity was retained in the mutant form of the protein. This is in agreement with the hypomorphic nature of *tni*, where the wild-type *TNI*transcript is also detected. In *tni* mutant, the catalytic inactivity of TNI^intron^ towards α and iso-linked poly-ubiquitin chains resulted in an inefficient turn-over of poly-ubiquitin into mono-ubiquitin. Since the two transcripts corresponding to wild-type TNI and aberrant TNI^intron^were detected almost in equal abundance in the mutant, it is likely that the normal and the aberrant forms of the TNI protein are expressed in comparable amounts considering their translation efficiency is similar. Thus, the total TNI protein in the *tni*cells comprises of TNI and TNI^intron^ which are indistinguishable immunologically due to their minor difference in molecular weight by ∼4 kDa. Recently, it was shown that the *da3-1* allele of *UBP14* had a G→T transversion at the 5′ exon-intron boundary of the last intron, generating a premature stop codon (Xu et al., 2016). Hence, the protein encoded by the *da3-1* locus was catalytically inactive since it lacked the C-terminal *Histidine*box essential for catalysis. Here, we show that UBP14 with its ZnF domain disrupted by an insertion of 34 amino acid residues (TNI^intron^) is also catalytically inactive. The ZnF-UBP domain from IsopeptidaseT (IsoT), the human orthologue of UBP14, binds to the di-glycine motif of the proximal ubiquitin in an un-anchored poly-ubiquitin chain and is essential for its catalytic activation (Reyes-Turcu et al., 2006). The ZnF-UBP domain of UBP14 shares 47% identity and 62% similarity with IsoT at the protein level (www.genome.jp/tools/clustalw/). Hence, it will be interesting to address whether such substrate-assisted catalysis is also conserved in UBP14. The catalytic inactivity of TNI^intron^led us to speculate that the 34-residue insertion in the ZnF domain of TNI^intron^ affects the substrate binding by TNI^intron^, rendering it catalytically inactive. Our data suggests that the full-length TNI^intron^ binds to Lys48-linked tetra-ubiquitin efficiently, implying that the retention of the intron-encoded 34 residues in TNI^intron^ does not abrogate its substrate binding. Moreover, ubiquitin binding by ZnF domain alone from the wild-type TNI and the mutant TNI^intron^ was comparable, indicating that the catalytic inactivity of TNI^intron^ was not due to inefficient substrate binding. Since the catalytic residues in the *Cysteine* and *Histidine* boxes were intact in TNI^intron^, an overall conformational change in TNI^intron^or steric hindrance by the 34-residue insertion perhaps rendered it inactive.

Based on the intact substrate binding by TNI^intron^, it can be argued that the mutant protein sequesters a portion of the *in vivo* substrate of TNI, making them unavailable to the functional TNI protein in the*tni* mutant. Thus, the gain-of-function nature of TNI^intron^ might have resulted in the*tni* phenotype. However, the heterozygous *tni* resembles wild-type, which understates the gain-of-function nature of *tni* allele. In the*tni* mutant, there was an overall increase in total ubiquitin compared to the wild-type. In eukaryotes, there exists a dynamic equilibrium between mono-ubiquitin and ubiquitin conjugates within a cell and any perturbation that affects this equilibrium results in an up-regulation of ubiquitin biosynthetic genes or acceleration of poly-ubiquitin to mono-ubiquitin formation by de-ubiquitinases (Park and Ryu, 2014). This contributes to the maintenance of cellular ubiquitin pool. Thus the reduced level of mono-ubiquitin in *tni* might have resulted in an increased poly-ubiquitin biosynthesis, accounting for the total increase in ubiquitin. Indeed, in the microarray data set, the poly-ubiquitin biosynthetic genes, *UBQ13* and *UBQ14* were 2-fold up-regulated in the *tni* mutant (Karidas et al., 2015).

TNI/ UBP14can hydrolyze both α-linked and iso-linked poly-ubiquitins (Doelling et al., 2001) whereas TNI^intron^is catalytically inactive. Thus,the α-linked poly-ubiquitin generated from ubiquitin translation as poly-proteins (Callis, 2014) and the iso-linked poly-ubiquitin generated during target protein degradation by 26S proteasome (Hershko and Ciechanover, 1992) contribute to the total increase in free poly-ubiquitin chains. The functional homologues of UBP14, *Sc*UBP14 from budding yeast and IsoT from human, have been shown to hydrolyze un-anchored poly-ubiquitin into mono-ubiquitin and thereby facilitate the degradation of proteins by 26S proteasome by preventing the accumulation of free poly-ubiquitin chains (Amerik et al., 1997; Wilkinson et al., 1995; Dayal, et al., 2009). Free Lys48-linked poly-ubiquitin chains can act as competitive inhibitors of 26S proteasome and thereby prevent substrate recognition and degradation by 26S proteasome (Amerik et al., 1997; Piotrowski et al., 1997). We observed that *tni* has higher abundance of Lys48-linked ubiquitinated proteins, suggesting an inefficient turn-over of proteins by 26S proteasome. Moreover, the *tni* mutant showed an increased response to MG132 treatment interms of its poly-ubiquitin profile, implying that the turn-over of target proteins by 26S proteasome is affected in *tni*. Mutations in the 19S regulatory subunits of 26S proteasome, *rpn10-1* and *rpn12-1* also result in higher levels of ubiquitin conjugates similar to that in *tni* mutant and exhibits reduced sensitivity towards exogenous auxin (Smalle et al., 2002, 2003). To examine whether *TNI* interacts with proteasomal subunit genes genetically, we crossed *tni* with regulatory particle subunit mutant *rpt2*(Lee et al., 2011). But we failed to establish double homozygous mutants(data not shown),possibly because poly-ubiquitin disassembly is largely affected when *tni* mutation is combined with that of regulatory particle subunits resulting in lethality of the double mutant. This further emphasizes the importance of *TNI* in protein turn-over by 26S proteasome.

The auxin-related growth defects of *tni*can largely, if not exclusively, be explained by the stabilization of some of the AUX/IAA transcriptional repressors. Most of the phenotypes of the *tni* mutant resembled those found in the gain-of-function mutations of AUX/IAAs where their stability is increased. The rootless phenotype of *tni* is similar to that of *bdl/IAA2*gain-of-functionand *mp/arf5*loss-of-function mutants (Hamann et al., 1999, 2002). *IAA12* works downstream to*ARF5*in hypophysis specification during embryogenesis. It has been shown that the stabilized form of IAA12 in the *bdl*mutant blocks auxin-dependent, ARF5-mediated activation of *TMO5*(*TARGET OF MP 5*) and *TMO7* required for hypophysis specification and root initiation (Schlereth et al, 2010). Venation patterning defects were also reported in *mp* and *bdl* mutants (Berleth et al., 2000). The severity of venation patterning defects in *tni* cotyledons was less than in*mp* and *bdl*, perhaps due to the hypomorphic nature of the *tni* allele. The reduced complexity of venation in *tni*cotyledons could be either due to a reduced auxin transport or signaling. *MP* expression is induced by auxin which in-turn regulates *PIN1* expression. Thus,*MP*-mediated activation of *PIN1* is required for venation patterning (Wenzal et al., 2007). However, we did not notice any significant change in *PIN1::GUS* expression between Col-0 and *tni* seedlings (Supplemental Fig. S7). Hence, the vascular patterning defects in *tni* might be due to the stabilization of AUX/IAAs resulting in reduced auxin signaling.

Fewer lateral roots in*tni* could be due to the stabilization of multiple AUX/IAAs including IAA3, IAA14, IAA18, IAA19 and IAA28 (Tian and Reed, 1999; Fukaki et al., 2002; Uehara et al., 2008; Tatematsu et al., 2004; Rogg and Bartel, 2001). Among the gain-of-function mutants isolated in these *AUX/IAAs*, *iaa14/slr*totally lacks lateral roots whereas the remaining mutants produce fewer lateral roots, suggestingthat these genes inhibit lateral root initiation. *ARF7* and *ARF19* act downstream to *IAA14/SLR*and activate the auxin responsive genes *LBD16* and *LBD29*which are required for lateral root initiation (Fukaki et al, 2002; Okushima et al, 2005, 2007). Genetic interaction of *tni* with *arf7-1* showed an additive effect in lateral root emergence, suggesting that both these genes contribute to the auxin-mediated lateral root development. It is possible that 26S proteasome-mediated degradation of IAA14 is further compromised in*iaa14tni* plants compared to the *iaa14* mutant alone, resulting in an additive phenotype. Moreover, the hypomorphic nature of *tni* allele might have also resulted in the enhanced phenotype, wheneverthe function of auxin signalingis abrogated as observed in *arf7-1 tni* and *tir1-1 tni*plants or auxin transport mutant *aux1-7*.

The dominant mutant *shy2-2* has altered sensitivity towards auxin and displays defects in lateral root formation, gravitropism and leaf morphology (Tian and Reed 1999). In *shy2-2*mutant, auxin response is attenuated due to the stabilization of IAA3/SHY2. We were unsuccessful in establishing the *shy2-2 tni*double homozygous mutant (data not shown). Stabilization of IAA18:GUS in various organs of *tni* could explain the reduced auxin sensitivity ofthe *tni* mutant. IAA18, along with IAA14, represses lateral root formation by sequestering ARF7 and ARF19 (Uehara et al., 2008). Hence the reduced lateral root formation in *tni* could be an additive effect of stabilization of both IAA14 and IAA18. In Arabidopsis, there are 29 AUX/IAAswith conserved domain II imparting stability to the protein (Overvoorde et al, 2005; Reed, 2001). AUX/IAAs interact with TIR1 through domain II and undergo poly-ubiquitination and subsequent degradation by 26S proteasome (Gray et al., 2001; Reed, 2001). All these AUX/IAAs follow a similar degradation mechanism by 26S proteasome with varying kinetics. Stabilization of DII:VENUS in *tni* is in agreement with the stabilization of multiple AUX/IAAs which resulted in reduced auxin response. We showedthat IAA18 and AXR3-NT are stabilized in *tni*. Stabilization of multiple AUX/IAAs in *tni* plants could provide an explanation for multiple auxin-related growth defects. Since *tni* heterozygous plants phenotypically resemble wild-type, it is unlikely that a dominant effect of the aberrant TNI^intron^ contributes to the stabilization of AUX/IAAs by sequestering them from TNI protein. A distinct possibility is that, the accumulation of poly-ubiquitin chains in *tni* creates a road-block for the degradation of AUX/IAAs by 26S proteasome. Similar proteasomal inhibition by free Lys48-linked poly-ubiquitin was reported earlier in yeast and humans (Amerik et al., 1997; Wilkinson et al., 1995; Dayal, et al., 2009; Piotrowski et al., 1997).

Collectively, our finding has uncovered that ubiquitin homeostasis is essential for the turn-over of AUX/IAA transcriptional repressors and the TNI/ *At*UBP14 de-ubiquitinase is implicated in the regulation of the stability of these repressors. Thus, TNI maintains normal auxin response in Arabidopsis through turn-over of AUX/IAAs by 26S proteasome. However, it is likely that target stabilization in the *tni* mutant is not specific to the AUX/IAA repressors and other proteins with high turn-over are also stabilized in the mutant. A proteomic approach to identify all the target proteins stabilized in *tni* compared to wild type will be able to test this possibility.

## CONCLUSIONS

In summary, we have identified the *tni* locus and shown that the *TNI* gene encodes the UBIQUITIN SPECIFIC PROTEASE 14 enzyme. UBP14 activity is partly compromised in *tni*s ince the mutation causes inefficient splicing of the primary mutant transcript, resulting in an aberrant protein. This leads to the stabilization of certain AUX/IAA repressor proteins and widespread auxin-deficient phenotype. As a result of the mutation, the *tni* organs accumulate poly-ubiquitin chains and excess poly-ubiquitinated proteins due to reduced ubiquitin protease activity.

## MATERIALS AND METHODS

### Plant materials

Inour study,we have used*Arabidopsis thaliana* ecotypesCol-0 and L*er*as wild-types. The mutants and the transgenic lines used here were reported earlier. These were *DR5::GUS* (Ulmasov et al., 1997), *DR5::nYFP* (Mahonen et al., 2015),*IAA2::GUS* (Marchant, et al., 2002),*HS::AXR3-NT:GUS* (N9571),*HS:axr3-1-NT:GUS* (N9572) (Gray et al., 2001), *DII-VENUS* (N799173), *mDII-VENUS* (N799174) (Brunoud et al., 2012), *IAA18:GUS* (Ploense et al., 2009), *arf7-1* (CS24607) (Okushima et al, 2005), *aux1-7* (CS3074) (Swarup et al, 2004), *tir1-1* (CS3798) (Ruegger et al, 1998), *pin1-5* (Yamaguchi et al., 2014) (N69067) and *ttn6-4* (CS16079) (Tzafrir et al., 2002). Most of these lines were obtained from Nottingham Arabidopsis Stock Center (NASC, UK) and Arabidopsis Biological Resource Center (ABRC, USA).

### Plant growth condition and treatments

Seeds were surface sterilized and stratified in dark for 2 days at 4°C following which they were transferred to the growth chamber maintained under long day conditions of 16 hours light (120 μmole/m^2^s) and 8 hours dark at 22°C.

For 1-NAA (Sigma, USA) and NPA (Calbiochem, Germany) sensitivity assays, seedlings were first grown for 4 days in NAA/NPA free ½ MS (Sigma) medium supplemented with 1% sucrose (Sigma, USA) and 0.8% agar (Hi Media, India). This was followed by the transfer of seedlings to 1-NAA or NPA containing medium. All the plates were placed vertically inside the growth for additional 3 (NAA) or 5 (NPA) days. Photographs were taken at 7 or 9 DAG of growth and lateral roots were counted using differential interference contrast (DIC) microscope (Olympus). 7-day old seedlings were treated with MG132 (Sigma) for 16 hrs in a liquid medium containing ½ MS salt.

### Gravitropic assay

Gravitropic response assay was performed according to the protocol described earlier (Hobbie et al, 2000). Briefly, the ½ MS plates containing the seeds were kept vertically inside the growth chamber. 4 days after germination, the plates were rotated 90° to the initial position. ImageJ software (*rsbweb.nih.gov/ij/)* was used to measure the angle of curvature after 3 days of gravistimulation.

### Tissue clearing

Seedlings were kept in 70% ethanol for 24 hrs followed by incubation in lactic acid for 30 mins. Seedlings were mounted in lactic acid on a transparent glass slide and observed under DIC microscope.

### Embryo dissection

Fertilized ovules were scooped out from siliques of matured plants and placed on a glass slide containing Hoyer’s medium (HM) and chloral-hydrate:glycerol:water in 8:1:2 ratio. Images were acquired using a Zeiss Axio Imager (Zeiss, Germany) M1 microscope with DIC settings.

### Map based cloning

The *tni*plant was crossed to L*er* accessionand inF_2 generation,_*tni*looking plants were selected. Genomic DNA was extracted from the inflorescence of individual *tni* plants using Nucleon Phytopure kit (GE HealthCare). *Polymorph*(www.polymorph.weigelworld.org<u>) and*dCAPSFinder (*</u>http://helix.wustl.edu/dcaps/dcaps.html) softwares were used to design CAPS and dCAPS markers (Supplemental Table S1). The *tni* locus was narrowed down to the interval of 65 kb with the help of the markers and 509 *tni* mapping population. Among 21 genes in this interval, the candidate genes were amplified and cloned into pGEM-Teasy vector (Promega, USA) followed by sequencing.

### Generation of constructs and transgenics

*TNI* coding sequence was PCR amplified from cDNA using specific primers P1888 and P1889 (Supplemental Table 2). To generate the*TNI* over-expression construct *35S::TNI*,the coding sequence of *TNI*was cloned downstream to *35S CaMV* promoterin *pCAMBIA 1302*. The artificial microRNA (amiR) against *TNI* was made as per the protocol described earlier (http://wmd3.weigelworld.org/cgi-bin/webapp.cgi). Briefly, primers P1918, P1919, P1920 and P1921 were used to clone fragment I, II, III and IV respectively and subsequently cloned into pGEM-T Easy vector. The primers P2887 and P2888 were used to amplify *amiR-TNI* and cloned into pCAMBIA1302. The *35S CaMV* promoter of *pCAMBIA1302* was replaced with RPS5a promoter to generate *RPS5a*::*amiR-TNI*(*amiR-TNI*). The *RPS5a* promoter sequence was amplified with P2231 and P2238. The 35S CaMV promoter was replaced with 1.5kb *pPLT7* in pCAMBIA1302 and pCAMBIA1390 using the primers P2801 and P2802. The CDS of *TNI^intron^* was cloned downstream to the *pPLT7* in *pCAMBIA1302* using NcoI restriction enzyme to make the construct *pPLT7::TNI^intron^*. Similarly *PLT7::amiR-TNI* construct was made.

To make the transcriptional fusion of *TNI*, 1.9 kb upstream region of *TNI* genomic region was amplified using primers P2759 and P2760 and cloned into pDONR221. The promoter sequence was cloned into pMDC162 destination vector to make *pTNI::GUS* construct.

All these constructs were transformed into *Agrobacterium* GV3101 by electroporation. Flowering Arabidopsis plants were transformed with *Agrobacterium* containing individual constructs using floral dip method (Clough and Bent, 1998).

*TNI* CDS was cloned in pGEX-4T-1 using primers P1912 and P1913 with engineered BamH1 and SalI restriction sites respectively to create *pGEX4-T-1-TNI* construct. *TNI*^intron^ of amplicon size 800 bp was amplified from *tni* cDNA with the primers P1888 and P1774 having BglII and BsmI restriction sites and cloned into *pGEM-T-TNI* to make *pGEM-T-TNI^intron^*. *pGEX4T-1-TNI* was replaced with *TNI^intron^*using SacI and HindIII restriction enzymes to make *pGEX4T-1-TNI^intron^* construct.

*TNI^C317S^* active site mutant was generated using Q5 Site-Directed Mutagenesis Kit (NEB).*pGEM-T-TNI* served as a template for making the site directed mutant using primers P2346 and P2347. *TNI* fragment in *pGEM-T-TNI* was replaced with *TNI^C317S^* using BglII and SpeI restriction enzymes to create *pGEM-T-TNI^C317S^*. From this vector, the *TNI^C317S^* coding sequence was moved to *pGEX4-T-1* using BamH1 and SalI restriction enzymes for protein expression.

The*pGEX-4T-1-TNI* was used as a template to generate various mutants; ZnF, ΔZnF, ΔUBA, ΔUBA2 and UBA domains using primer pairs P1912, P2405; P2403, P1913; P1912, P2549; P1912, P2806 and P2827, P2808 respectively with engineered BamH1 and Sal1 restriction sites. All these deletion constructs were subsequently cloned into pGEX-4T-1.

### Purification of GST tagged recombinant proteins from *E.coli*

BL21 *E.coli* strain was used for recombinant protein expression. *E.coli*cells were induced with 0.5 mM IPTG (Sigma, USA) at mid-log phase and incubated at 16°C for 12 hrs. Cells were harvested by centrifugation at 5000 rpm for 5 mins at 4°C. The cell pellet was suspended in cell lysis buffer (50 mM Tris-HCl pH 7.4, 150 mM KCl, 0.5 mM EDTA, 10% glycerol, 0.5% NP-40, 1 mM PMSF and protease inhibitor cocktail)followed by sonicationuntil it turned clear. The clear supernatant was collected after centrifugation at 12,000 rpm for 20 mins at 4°C. The supernatant was incubated with glutathione beads (Novagen, USA) for 2 hrs at 4°C with constant shaking. The beads were washed five times with ice-cold cell suspension buffer. 2 mM glutathione was used to elute the bound proteins. SDS-PAGE was run to check the purity of recombinant proteins.

### *In vitro* and *in vivo* de-ubiquitination assay

*In vitro*and *in vivo*de-ubiquitination assays wereperformed as mentioned earlier (Turcu et al, 2006; Rao-Naik et al., 2000). Briefly, an assay reaction of 50μl comprising equal concentration of purified protein, 50 mM Tris-HCL pH 8, 150 mM Nacl, 10 mM β-mercaptoethanol, 0.5 mg/ml BSA and 2μg 2-7 mer of poly-ubiquitin (Boston Biochemical) was incubated at 37°C for 3 hours. Lamelli buffer was added to stop the reaction. The assay mixture was fractionated by SDS-PAGE and immunoblot analysis was done using anti-ubiquitin antibody (Novus Biologicals, USA). For *in vivo* assay, construct was co-transformed in *E.coli*along with the p8190-UBQ10 and pACYC184-Ub_4_(substrates) (Rao-Naik et al., 2000). *E.coli*cells were induced with 0.5 mM IPTG and incubated for 3 hours at 37°C. Equal number of cells was pellet down and suspended into cell lysis buffer. Cell suspension was subjected to sonication. The supernatant was boiled with Lamelli buffer and run into SDS-PAGE. Immuno-blotting was performed using anti-ubiquitin antibody and anti-His (Sigma, USA).

### *In vitro* substrate binding assay

The GST-tagged recombinant proteins were purified using glutathione sepharose beads as mentioned earlier. 5-50 μl of bead bound protein was incubated with 1μg Lys48 linked tetra-ubiquitin (Boston Biochemicals, USA) in sodium phosphate, pH 7.4 in presence of protease inhibitor cocktail (Roche) (Medina et al., 2012). The reaction mixture was kept for incubation at 4°C for 2 hrs with constant shaking. Lamelli buffer was added to stop the reaction and the sample was boiled for 10 mins. The reaction mixture was fractionated by SDS-PAGE and immune-blotting was performed using anti-ubiquitin antibody.

### GUS staining and MUG assay

GUS staining was performed as described earlier (Karidas et al., 2015). Briefly, seedlings were harvested in ice-cold 90% acetone and kept on ice for 20 mins. Seedlings were then incubated in staining buffer (0.5 M sodium phosphate buffer pH 7.2, 10% Triton-X, 100 mM potassium ferrocyanide and 100 mM potassium ferricyanide) for 20 mins at room temperature. 2 mM X-Gluc (Thermo Scientific) was added to the fresh staining buffer containing seedlings and incubated at 37°C for 12 hours. The seedlings were washed with 70% ethanol for 30 mins to 1 hour at RT until the chlorophyll had been removed. The tissues were mounted on a glass slide in lactic acid and observed under DIC microscope.

MUG assay was performed according to the protocol mentioned earlier (Weigel and Glazebrook 2002). Briefly, proteins were extracted from the seedlings using extraction buffer (50 mM sodium phosphate pH 7.0, 10 mM EDTA, 0.1% SDS, 0.1%TritonX-100, 1 mM PMSF and protease inhibitor cocktail). Protein extract was centrifugedat 12,000 rpm at 4°C for 15 min. The clear supernatant was collected and equal concentration of proteins wastaken for the MUG assay. The MUG assay buffer is same as that of the extraction buffer supplemented with 1 mM MUG (Sigma). The reaction was incubated at 37°C water bath for 20 min. The reaction was stopped using 0.2 M sodium carbonate. TECAN fluorimeter was used to measure the fluorescence at 365nm excitation and 455nm emission wavelength.

### Antibody generation

The anti-TNI polyclonal antibody was raised against a synthetic peptide (USV Ltd, India) corresponding to 156-174 residues of TNI/ UBP14, which was identified an a potent immunogen using the software http://tools.iedb.org/main/bcell/. 5 mg of peptide conjugated with KLH carrier, with a purity of greater than 85%was synthesized. Rabbits were first immunized with 1mg of conjugated peptide, followed by 3 booster immunizations, each with500 µg of conjugated peptide. After immunizing, the post immune sera was collected from rabbit.

### Immunoblot analysis

Proteins were isolated from 7-8-day old seedlings using protein extraction buffer (50 mM Tris-HCl (pH 7.4), 300 mM KCl, 0.5 mM EDTA, 10% glycerol, 0.5% NP-40) along with 1 mM PMSF, 50μM MG132 and complete protease inhibitor cocktail (Roche). The protein extracts were cleared by centrifugation at 12,000 rpm for 15 mins at 4°C. Equal concentrations of protein were run in 15% and 10% SDS-PAGE. After electrophoresis, proteins were transferred to the PVDF membrane (Millipore) and immunoblot analysis was performed using anti-GFP (Roche), anti-ubiquitin, anti-Lys48 ubiquitin (CST) and anti-Lys63 (Enzo Life sciences) antibodies. ECl (Millipore) was used to develop the blot in chemidoc.

### Confocal microscopy

Roots were stained with propidium iodide (10μg/mL) (Sigma) for 5 mins and mounted in a glass slide. Roots were observed under the laser confocal microscope (Zeiss LSM 710).

### RNA isolation and cDNA synthesis

Total RNA was extracted from 7-day old seedlings using Trizol (Sigma). The extracted RNA was treated with DNase (Fermentus) for 2 hrs at 37°C and precipitated by sodium acetate. 2μg of RNA was taken as a template for cDNA preparation using Revert Aid M-MuLV reverse transcriptase (Fermentus). A 20 μl reverse transcription reaction was set up and the amplified product was visualized using ethidium bromide stained 1% agarose gel.

## ACCESSION NUMBERS

The accession numbers of the genes mentioned in this article are given below and their sequence data can be found in Arabidopsis genome initiative (WWW.ARABIDOPSIS.ORG):*IAA2* (*At3G23030*), *AXR3* (*At1G04250*), *IAA18* (*At1G51950*), *ARF7* (*At5G20730*), *AUX1* (*At2G38120*), *TIR1* (*At3G62980*), *PIN1 (At1G73590*) and *TTN6* (*At3G20630*).

## ACKNOWLEDGEMENTS

We acknowledgeB. Vijayalakshmi Vadde (Indian Institute of Science, Bangalore, India) for raising the antibody against *TNI*, Sowmya Spandana M (Indian Institute of Science, Bangalore, India) for help in making clones, Bonnie Bartel (Rice University, Texas, US) for *pIAA28::IAA28:myc* line, Jason Reed (University of North Carolina, Chapel Hill, US) for *IAA18:GUS*, Judy Callis (University of California, Davis, US) for *ScUBP14*, *Ub_6_* and *His-Ub_4_* constructs, Ben Scheres (Wageningen University, Netherlands) for *DR5::GUS* and *DR5::nYFP* lines and Kalika Prasad (IISER Thiruvananthapuram, India) for *pPLT7* construct.

