## Supplemental data for "The TARANI/ UBIQUITIN SPECIFIC PROTEASE 14 destabilizes the AUX/IAA transcriptional repressors and regulates auxin response in *Arabidopsis thaliana*"

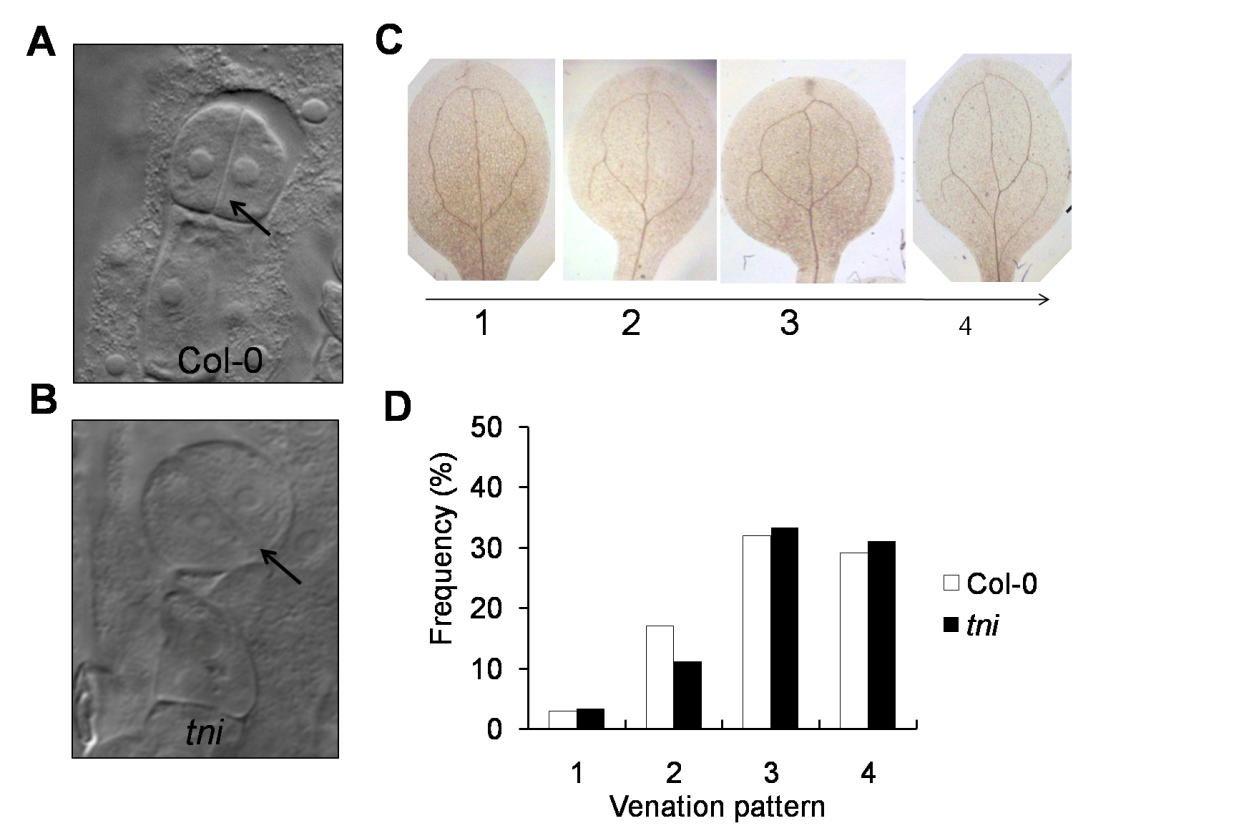

**Supplemental Figure S1. (A) and (B)** 2-cell pro-embryos. **(A)** The vertical cell division pattern in all the apical cell of Col-0 (N=30) and oblique pattern of cell division **(B)**in 8% *tni*pro-embryo (N=87) indicated by black arrows. **(C)** The cleared cotyledons of Col-0 in increasing order of vascular complexity indicated by numbers. **(D)** The frequency distribution of Col-0 and *tni*cotyledons with indicated venation pattern. N=103 (Col-0) and 90 (*tni*).

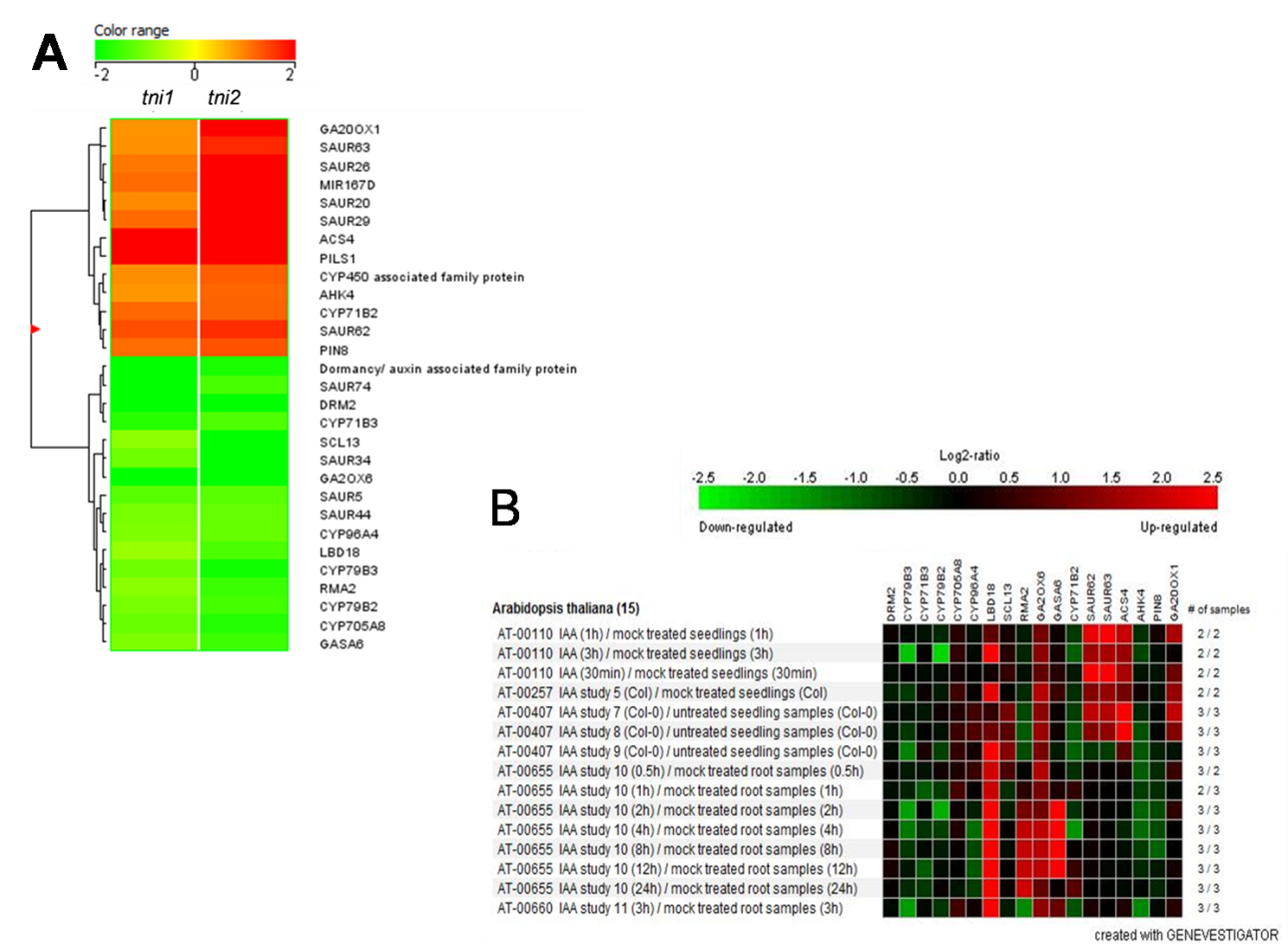

**Supplemental Figure S2.Differentially expressed auxin-related genes in *tni*mutant.(A)** Heat-map of the microarray data showed the differentially expressed auxin-related genes in two biological replicates of *tni*. **(B)**Genevestigator database search showed the expression of some of the differentially expressed auxin-related gene set in *tni***(A)** are also perturbed in exogenously auxin treated seedlings.

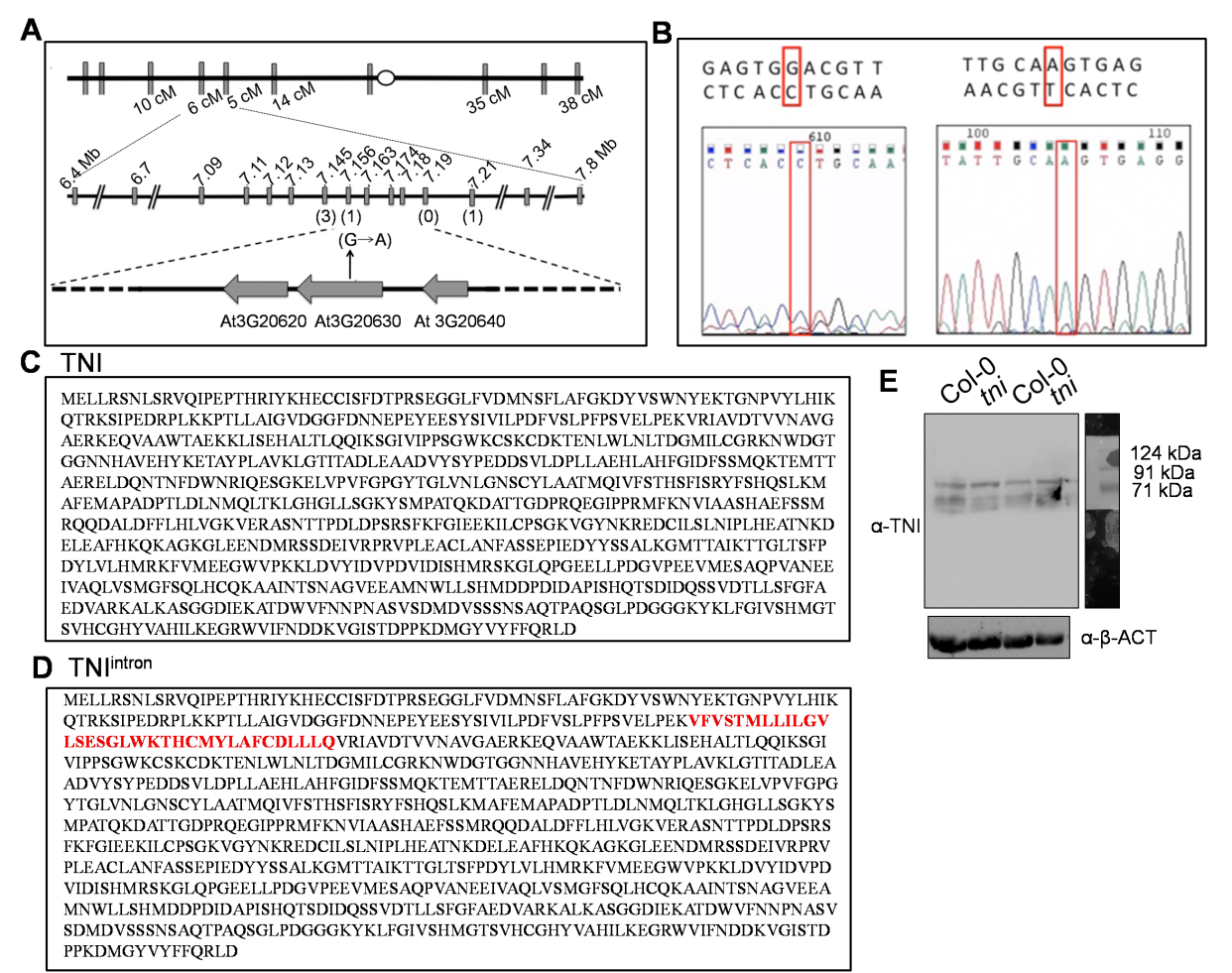

**Supplemental Figure S3. Fine mapping of *tni* locus and TNI protein sequence (A)** Schematic of the 3^rd^ chromosome (bold grey line) shows molecular markers (vertical grey bars) and their respective recombination frequencies. The numbers above the markers at the lower panel indicate their physical position in megabases (Mbs). Numbers in parenthesis indicate the number of recombinants obtained at that marker. The grey block arrows denote annotated genes in that locus. The black arrow shows G to A transition in *At*3G20630 which codes for *UBP14.***(B)** Sequencing chromatograms of Col-0 (left) and *tni* (right). The blue and red boxes denote sequence and chromatogram respectively show the transition from G to A in *tni* denoted by red box**(C)** The amino acid sequence of full length TNI (797 amino acids) and TNI^intron^ (831 amino acids) **(D)**. The intron 3 encoded 34 amino acid sequence is denoted in red **(D)**which is in-frame to the TNI coding sequence.**(E)** Full Western blot image (shown in Figure 5J) of the total protein extracted from two biological replicates of Col-0 and *tni* seedlings probed with anti-TNI antibody. Anti-β-ACT was used as loading control.

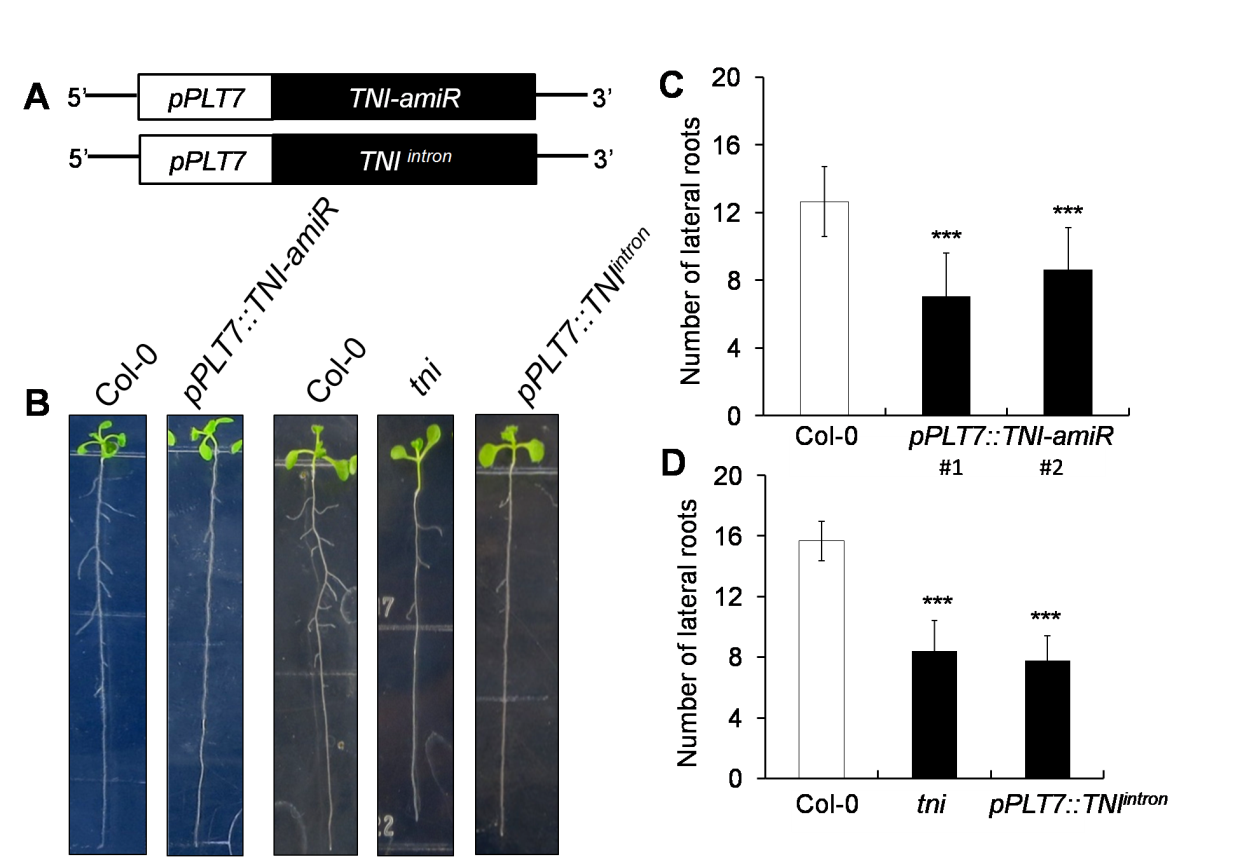

**Supplemental Figure S4. Effects of mis-expression of *TNI^intron^* and down-regulation of *TNI* transcript on the lateral root formation. (A)** The schematic representations of the constructs used to down-regulate *TNI* transcript and mis-express *TNI^intron^***)** in lateral root specific promoter, *PLT7* promoter (*pPLT7*). (B) Seedlings highlighting lateral roots in the transgenics and the controls. **(C)**and**(D)** The average number of lateral roots of 11-day old seedlings. Error bars represent SD. Statistical analysis was done by unpaired Student’s *t*-test. *** denotes p<0.0001. N=11(Col-0); 22 (#1); 16(#2) in **(C).** N=12(Col-0); 10(*tni*); 28 (*pPLT7*::*TNI^intron^*) in **(D)**

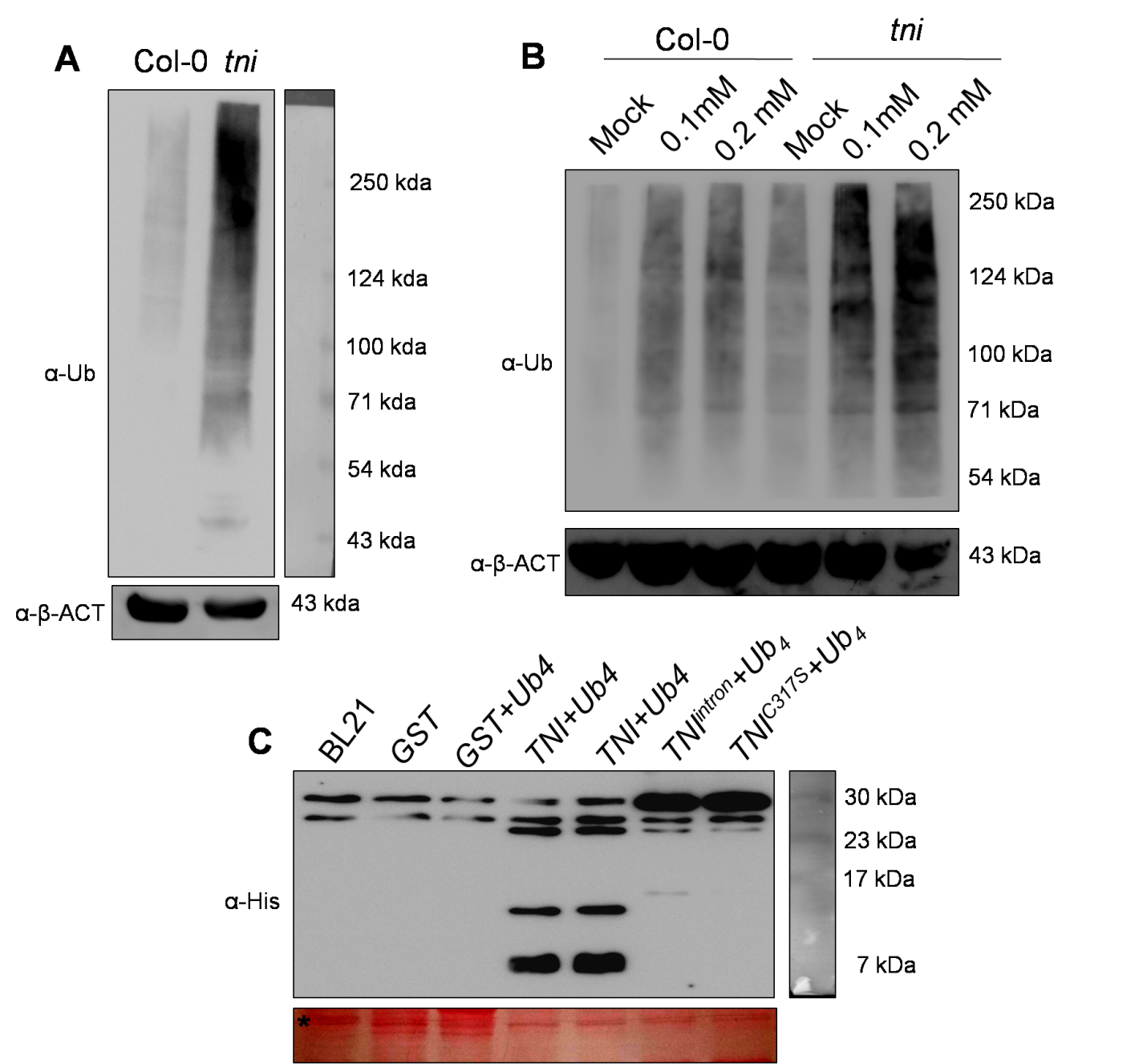

**Supplemental Figure S5. Poly-ubiquitinated protein in *tni* mutant (A)**Anα-Ub Western blot of total proteins extracted from 7-day old seedlings and resolved in a 10% SDS-PAGE. Exorbitant accumulation of poly-ubiquitinated proteins (appear as smear) in *tni* relative to Col-0. The α-β-ACT Western blot served as loading control. Numbers in the right indicate molecular weight of the protein marker. **(B)** Western blot of total proteins extracted from 7-day old seedlings treated with 0 (Mock), 0.1 or 0.2 mM MG132 for 16 hrs. The proteins are resolved in a 10% SDS-PAGE. Smears indicate poly-ubiquitinated target proteins. Anti-β-ACT Western blot served as a loading control. Molecular weight of the protein markers are shown on the right. (**C)**An α-His Western blot of *E. coli* lysates expressing N-terminal His-tagged tetra-ubiquitin chain (Ub_4_) along with recombinant GST-TNI, GST-TNI^intron^ and GST-TNI^C317S^ fusion proteins. Lysates expressing no recombinant proteins (BL21), recombinant GST alone (GST) and recombinant GST plus Ub_4_were used as negative controls. Numbers on the right indicate molecular weights of the marker protein. A Ponceau-stained membrane shown below served as loading control.

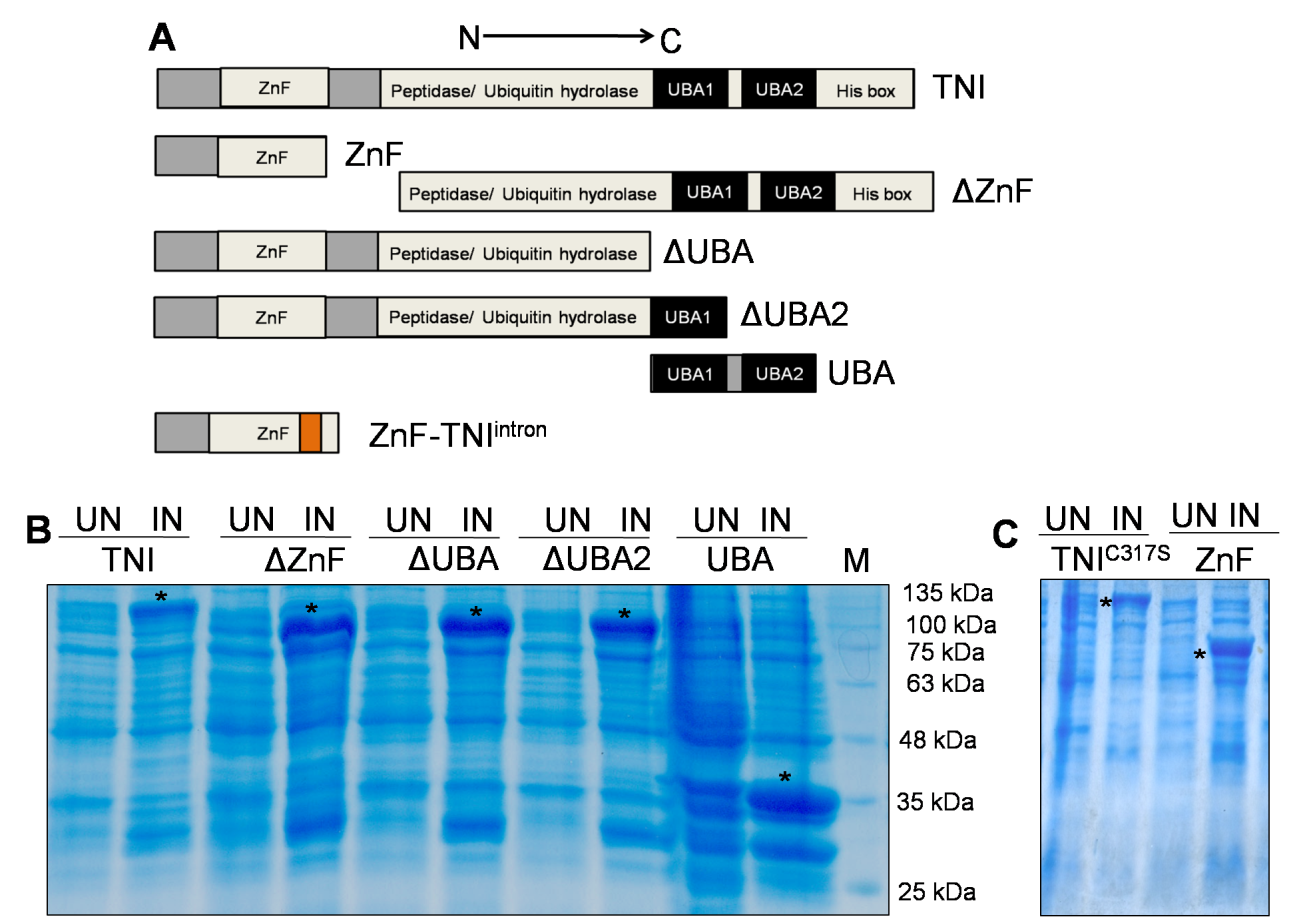

**Supplemental Figure S6. (A)**The domain architectures of TNI and its deletion forms. **(B)**and **(C)** Coomassie-stained poly-acrylamide gel highlighting the bands corresponding to GST-tagged full length TNI, 114 kDa; ΔZnF, 83 kDa; ΔUBA, 91 kDa; ΔUBA2, 97 kDa; UBA, 34 kDa, TNI^C317^, 114 kDa and ZnF, 60 kDa. Asteriks denotes the induced protein band. UN, uninduced; IN, induced and M, protein molecular weight marker.

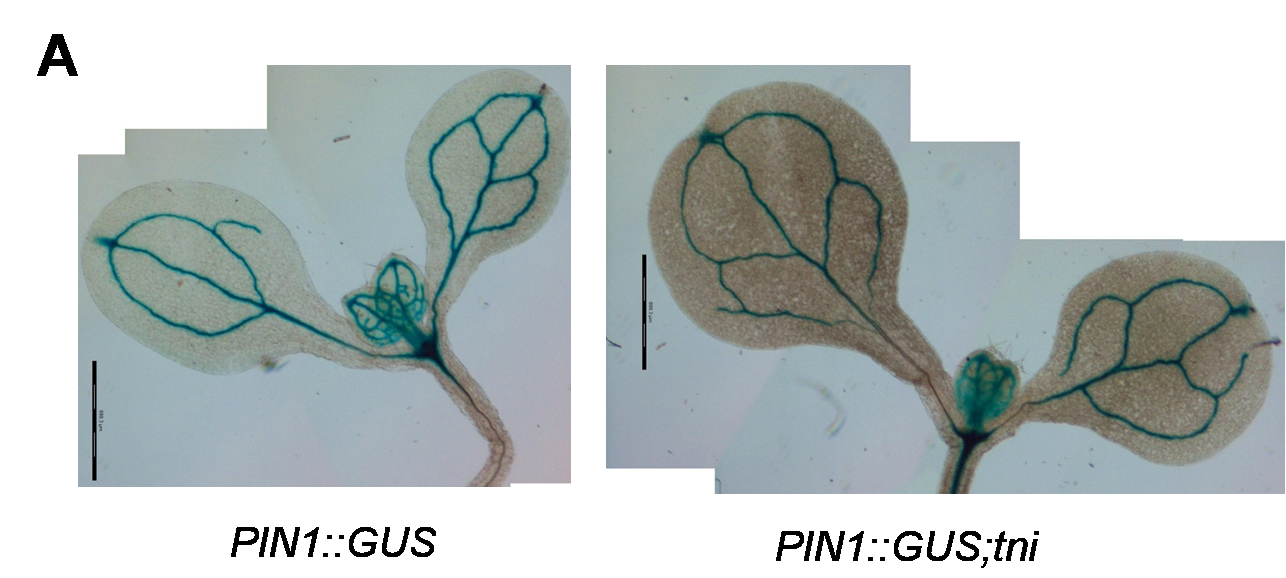

**SupplementalFigure S7.***PIN1::GUS* expression in 6-day old wild-type (Ler) and PIN1::GUS;*tni*seedlings.

**Supplemental Table S1.**List of markers used for fine mapping of *TNI* locus with their nature and positions in the genome.
